# ARHGAP20 organizes spatial Rap1–RhoA signaling coordination controlling adhesion dynamics during migration

**DOI:** 10.64898/2026.07.19.739419

**Authors:** Sandra Pagano, Colline Sanchez, Natasha Cox-Cammer, Ravi M. Bhalla, Roshan Ravishankar, Deisy Segura-Villalobos, Anne Muesch, Julio A. Aguirre-Ghiso, Gaudenz Danuser, Louis Hodgson

**Author notes:** Authors contributed equally (Sandra Pagano, Colline Sanchez). Correspondence should be addressed to: Louis Hodgson, PhD.

## Abstract

Cell migration requires the precise coordination of signaling pathways that regulate cytoskeletal dynamics and adhesion turnover. Rho GTPase–activating proteins (RhoGAPs) play critical roles in shaping these processes by controlling the spatial and temporal activity of small GTPases. ARHGAP20 is a RhoA-specific GAP, a downstream target of the Ras-related GTPase Rap1, and has been implicated in cancer cell motility, yet its functional role in coordinating migration-associated signaling remains poorly understood. Here, we investigated the role of ARHGAP20 in cell migration and its impact on the coordination between adhesion- and contractility-associated signaling pathways regulated by Rap1A and RhoA respectively. Using loss-of-function approaches in MTLn3 cells, we show that depletion of ARHGAP20 impairs both directed and random migration, leading to reduced cell velocity and displacement, and increased cell adhesion.

To explore the underlying signaling mechanisms, we developed a genetically encoded FRET biosensor to monitor Rap1A activity and combined it with a near-infrared RhoA biosensor to simultaneously analyze their spatiotemporal dynamics in living cells. We found that Rap1A and RhoA activities are negatively coordinated during leading-edge dynamics and that ARHGAP20 depletion enhances this local anticorrelation. To further define where ARHGAP20 regulates Rap1A–RhoA signaling, we applied a microdomain-based analytical framework to quantify local signaling clusters. This analysis revealed that ARHGAP20 selectively modulates Rap1A–RhoA coordination outside focal adhesion regions, while leaving signaling correlations and overlap within focal adhesions unchanged.

Subcellular localization analyses further reveal that ARHGAP20 is largely excluded from focal adhesions but associates with microtubules, the endoplasmic reticulum, the Golgi apparatus, and multiple Rab-positive vesicular compartments, supporting a trafficking-dependent mechanism for its spatial targeting. Together, our results identify ARHGAP20 as a regulator of cell migration that modulates the coordination between Rap1A and RhoA signaling through intracellular vesicular trafficking, highlighting the GAP’s role in organizing the spatial coupling between signaling pathways in adhesion-associated regions required for efficient cell migration.

## Introduction

Cell migration is a fundamental biological process that underlies numerous physiological and pathological events, including tissue morphogenesis (1–3), immune responses (4–7), wound healing (2, 3), and cancer dissemination (8–12). Efficient migration requires the coordinated regulation of actin cytoskeleton remodeling, membrane protrusion, and the dynamic formation and turnover of cell–matrix adhesions (7, 9, 13–21). These processes must be precisely organized in space and time to allow cells to generate protrusive forces at the leading edge, establish adhesive contacts with the extracellular matrix, and produce contractile forces that drive forward movement (10, 11, 22–24).

Small GTPases of the Ras superfamily play central roles in orchestrating these signaling networks (25–28). RhoA and Rap1 are two small GTPases playing key roles in cell motility and migration. Rho subfamily GTPases, including RhoA, regulate actomyosin contractility and cytoskeletal organization (27, 29–32), whereas Ras-related GTPases, including Rap1, contribute to the control of adhesion signaling and membrane trafficking (33–35). The activity of these molecular switches is tightly regulated by guanine nucleotide exchange factors (GEFs), which promote their activation, and GTPase-activating proteins (GAPs), which stimulate GTP hydrolysis and terminate signaling (13, 18, 21, 25, 28, 36). Through this regulatory network, GAP proteins play critical roles in shaping the spatial and temporal organization of small GTPase signaling during cell migration (18, 25, 37–40), but they remain not as well characterized as GEFs within the context of phenotypic outcomes through specific GTPase targeting.

ARHGAP20 is a RhoA-specific GAP that has been implicated in the regulation of cytoskeletal dynamics and cancer cell motility (41–43). Previous studies have suggested that ARHGAP20 acts downstream of the Ras-related GTPase Rap1 signaling to regulate RhoA activity, linking adhesion signaling to the control of actomyosin contractility (41–43). Consistent with this idea, ARHGAP20 expression has been reported to be elevated in early disseminating HER2+ mammary tumor cells, a population characterized by increased motility, invasive and systemic disseminating potential compared to later-stage metastatic cells (44). These observations suggest that ARHGAP20 may contribute to the regulation of signaling pathways that coordinate adhesion and contractility during cell migration. However, little is known about the subcellular mechanisms that spatially position ARHGAP20 to regulate localized RhoA signaling during migration.

Rap1 is a major regulator of integrin-mediated adhesion and promotes the formation and stabilization of adhesive structures at the leading edge of migrating cells (7, 39, 45, 46). In contrast, RhoA regulates actomyosin contractility and contributes to adhesion maturation and force generation (9, 14, 21, 36, 47, 48). Therefore, the dynamic balance, in space and time, between Rap1-dependent adhesion signaling and RhoA-mediated contractility is essential for efficient migration. Because ARHGAP20 acts as a RhoA-targeting GAP downstream of Rap1, it is well positioned to contribute to the coordination between these pathways. However, the mechanisms by which ARHGAP20 regulate the spatial and temporal organization of Rap1 and RhoA signaling during migration remain poorly understood. Although ARHGAP20 has been implicated in the regulation of cell motility, how it contributes to the coordination of signaling pathways controlling adhesion and contractility remains unclear. Because ARHGAP20 is positioned at the interface between Rap1- and RhoA-dependent signaling, it may play an important role in organizing the signaling transitions required for efficient migration. In addition, whether ARHGAP20 is statically associated with adhesion structures or instead reaches these signaling sites through intracellular trafficking has not been investigated. Among the two Rap1 isoforms, Rap1A has been particularly implicated in the regulation of adhesion signaling at the plasma membrane, including the formation of membrane protrusions and integrin-dependent adhesion complexes that drive migration (45, 49–52). This isoform-specific localization and function make Rap1A the most relevant Rap1 paralog for studying adhesion-associated signaling during cell migration.

Here, we examined how ARHGAP20 regulates the migratory behavior of rat mammary adenocarcinoma MTLn3 cells and the coordination of Rap1A and RhoA signaling during cell migration. Using quantitative migration assays together with live-cell multiplexed Förster resonance energy transfer (FRET) biosensor imaging, we first characterized the spatiotemporal coordination between Rap1A and RhoA activities during leading-edge dynamics. We then applied a microdomain-based analytical framework to quantitatively map local Rap1A–RhoA signaling coordination throughout the cell and identify the subcellular regions in which ARHGAP20 regulates this signaling network (53). Combined with quantitative analyses of ARHGAP20 subcellular localization, these studies reveal intracellular trafficking compartments as the principal sites of ARHGAP20 localization, providing a mechanistic framework for how this RhoGAP is spatially targeted to regulate signaling adjacent to, rather than within, focal adhesions. Together, our results identify ARHGAP20 as a regulator of signaling coordination between adhesion and contractility pathways during breast cancer cell migration.

## Results

### ARHGAP20 depletion impairs migration of MTLn3 cells

ARHGAP20 is upregulated by approximately fivefold in an RNA-seq study of early-stage disseminating HER2+ mammary tumor cells (44). These early evolved cancer cells exhibit elevated motility, invasive and systemic disseminating potential, consistent with alterations in gene expression that drive their characteristic phenotype. (44). To investigate the role of ARHGAP20 in cell migration, we performed a partial genetic depletion using siRNA in highly motile rat adenocarcinoma cell line MTLn3, with knockdown efficiency quantified by RT–qPCR. This approach resulted in an approximately 32% reduction in ARHGAP20 mRNA levels compared with control siRNA transfection (siCTL) (Fig. 1A). Given the previously reported role of ARHGAP20 in the regulation of cancer cell migration, we first examined the effect of ARHGAP20 depletion on directed migration using a Boyden chamber assay in the presence of a serum gradient. Depletion of ARHGAP20 significantly reduced the number of cells migrating toward the serum-enriched compartment compared with control conditions (Fig. 1B), indicating that ARHGAP20 promotes chemotactic migration in MTLn3 cells. To further dissect how ARHGAP20 contributes to migration, we performed random two-dimensional migration assays and tracked MTLn3 cells for 6 hours following transfection with either siCTL or siRNA targeting ARHGAP20 (siARHGAP20) (Fig. 1C). Trajectory analysis revealed that ARHGAP20 depletion significantly reduced both the mean migration distance (Fig. 1D) and the average cell velocity (Fig. 1E), indicating an overall impairment of cell motility. To visualize the motility of cells over time, individual cell trajectories were plotted relative to their point of origin. Under control conditions, most cells progressively moved away from their origin, reaching distances of up to ∼80 µm. In contrast, ARHGAP20-depleted cells remained largely confined near their starting position, with most cells remaining within ∼20 µm of their origin, although a small subset reached distances of 50 µm (Fig. 1F, G). To quantify this behavior, we measured the mean distance from the origin over time (Fig. 1H). Control cells rapidly moved away from their point of origin, followed by a slower phase of displacement while continuing to move outward, reaching an average distance of ∼40 µm after 6 hours. In contrast, ARHGAP20-depleted cells exhibited a reduced displacement and failed to sustain outward movement, resulting in a final average distance of only ∼20 µm from their origin. Consistently, quantification of the mean distance from the origin at the end of the experiment revealed an approximately twofold reduction in global displacement upon ARHGAP20 depletion (Fig. 1I). Together, these results demonstrate that ARHGAP20 plays an important role in regulating the migratory capacity of MTLn3 cells by maintaining cell velocity, displacement, and directional persistence.

**Fig 1.**
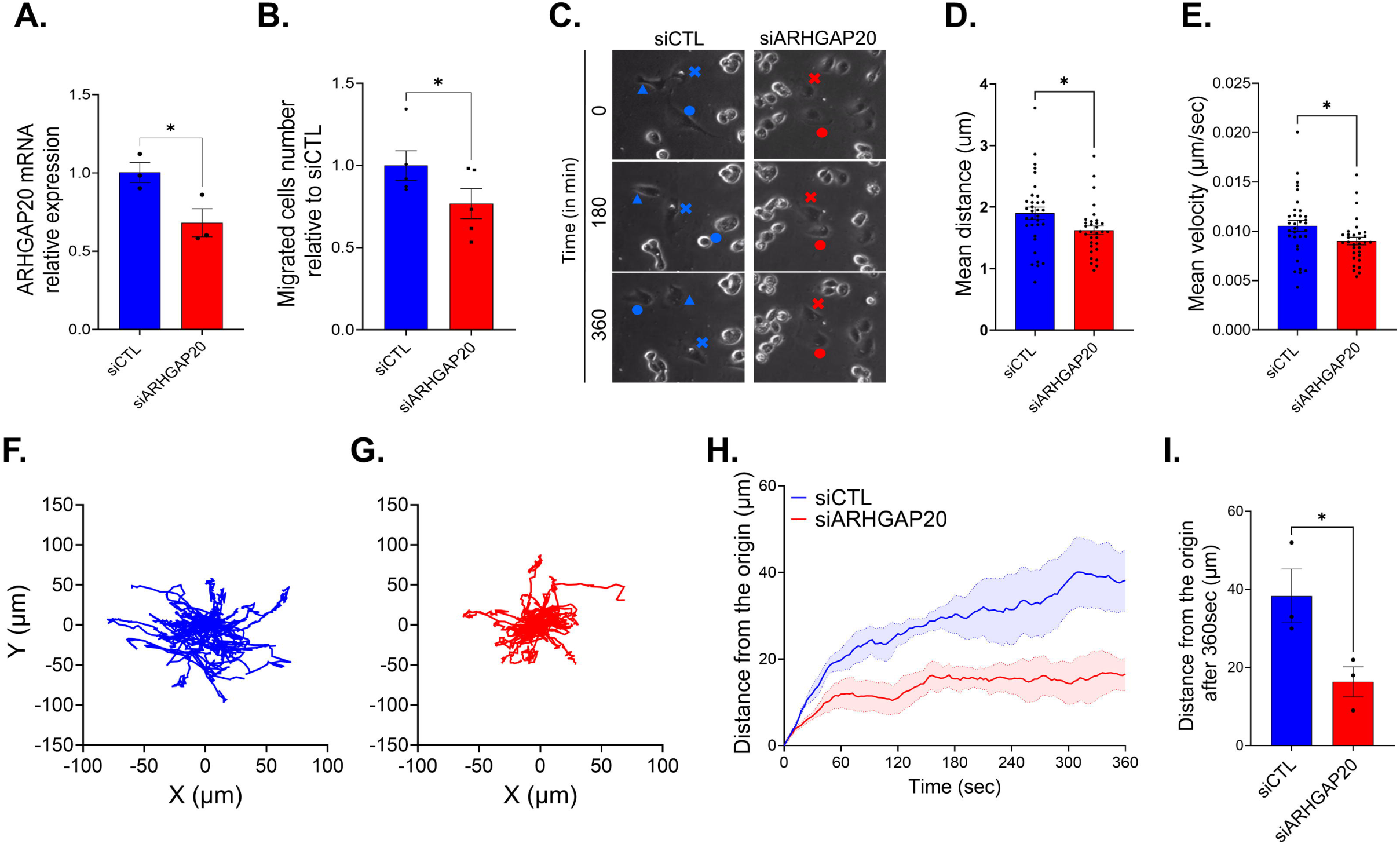
ARHGAP20 depletion impairs migration of MTLn3 cells. (A) Relative expression levels of ARHGAP20 in siCTL or siARHGAP20 transduced MTLn3 cells. Data are presented as mean ± standard error of the mean (SEM) from 3 independent experiments. (B) Directed migration assay performed using a Boyden chamber in the presence of a serum gradient in MTLn3 cells transfected with control siRNA (siCTL) or ARHGAP20 siRNA (siARHGAP20). Data represent mean ± SEM from 5 independent experiments (C-I) Random migration assays of MTLn3 cells. Cell movements were tracked for 6 h and trajectories were aligned to a common origin for visualization. (C) Representative trajectories of individual MTLn3 cells, each cell identified by a distinct symbol (arrow, cross, circle) to track its movement, in siCTL or siARHGAP20 conditions(D) Quantification of the mean migration distance traveled by individual cells. (E) Quantification of the mean cell velocity. (F–G) Spatial distribution of individual cell trajectories plotted relative to their point of origin for siCTL (F) and siARHGAP20 (G) conditions. (D-G) Each point represents a single tracked cell. (H) Mean distance from the origin plotted as a function of time for siCTL and siARHGAP20. (I) Quantification of the mean distance from the origin at the end of the experiment. (C-I) Data represent mean ± SEM from 3 independent experiments. Statistical significance was determined using an unpaired two-tailed Student’s t-test. *p<0.05

### Development and validation of a genetically encoded Rap1A FRET biosensor

To investigate how ARHGAP20 regulates cell migration, we next examined how the presence or absence of ARHGAP20 affects the coordination of activity between its upstream activator Rap1A and its downstream effector RhoA, two key regulators of cell motility (7, 9, 39, 45, 47, 54). To enable direct visualization of Rap1A activity in living cells, we developed a genetically encoded, single-chain Förster Resonance Energy Transfer (FRET) biosensor for Rap1A based on the cyan-yellow fluorescent proteins pair. The biosensor is composed of two monomeric fluorescent proteins, mCerulean3 (donor) and mVenus (acceptor), separated by a flexible linker of optimized length (55). Full-length Rap1A was positioned at the C-terminus of the biosensor to preserve its hypervariable region and the C-terminal CAAX motif required for proper membrane targeting. The N-terminus contains the Ras-binding domain (RBD) of Ral guanine nucleotide dissociation stimulator (RalGDS), which serves as a Rap1-specific binding domain (Fig. 2A). Upon Rap1A activation, intramolecular binding between GTP-bound Rap1A and the RalGDS-RBD changes the relative orientation of the fluorescent proteins, increasing FRET efficiency. Fluorescence emission spectra recorded following donor excitation revealed a clear difference between active and inactive forms of the biosensor. Cells expressing the constitutively active Rap1A mutant (G12V) displayed approximately 2-fold higher FRET compared with the inactive mutant (S17N), indicating a good dynamic range of the biosensor (Fig. 2B). We further extended this design to the Rap1B isoform and observed a similar fluorometric response (Fig. S4), indicating that the biosensor architecture is compatible with both Rap1 isoforms. To further characterize the biosensor, we analyzed the FRET ratio of the wild-type biosensor and several Rap1A mutants affecting nucleotide binding or effector interaction (Fig. 2C). Constitutively active mutants (G12V and Q63E), which impair GTP hydrolysis and stabilize the GTP-bound state, display FRET ratios comparable to the wild-type biosensor. This likely reflects the high level of biosensor expression in this experimental system, which results in a large fraction of Rap1A being maintained in an active state when cells are attached (Fig. 2D). In contrast, the inactive mutant S17N, which preferentially binds GDP, significantly reduced biosensor activity. Mutations affecting the effector-binding interface of Rap1A (D38A, D38E, T35S, and Y40C), which disrupt interaction with downstream effectors such as RalGDS, also markedly decreased the FRET signal (Fig. 2C). These results demonstrate that the biosensor signal depends on the interaction between GTP-bound Rap1A and the RalGDS effector domain. To further test the specificity of this interaction, we replaced the RalGDS Ras-binding domain with the p21-binding domain (PBD) of PAK, commonly used in pull-down assays to detect active Rac1 and Cdc42. Substitution of the RalGDS domain with the PBD strongly reduced the FRET signal, indicating that efficient biosensor activation requires the specific interaction between Rap1A and the RalGDS binding domain (Fig. 2C). One concern with biosensors of this design is the extent to which endogenous downstream effectors might compete with the intramolecular binding of the activated GTPase to its cognate binding domain within the biosensor backbone. We tested this by pull-down assays. An activated mutant form of the biosensor could be efficiently pulled down by the excess exogenous GST-RalGDS domain only when its cognate RalGDS binding domain was replaced with the PBD. These results indicate that, in the activated state, the GTPase within the biosensor backbone interacts predominantly with its cognate binding domain rather than with competing endogenous targets (Supplemental Fig. S1).

**Fig 2.**
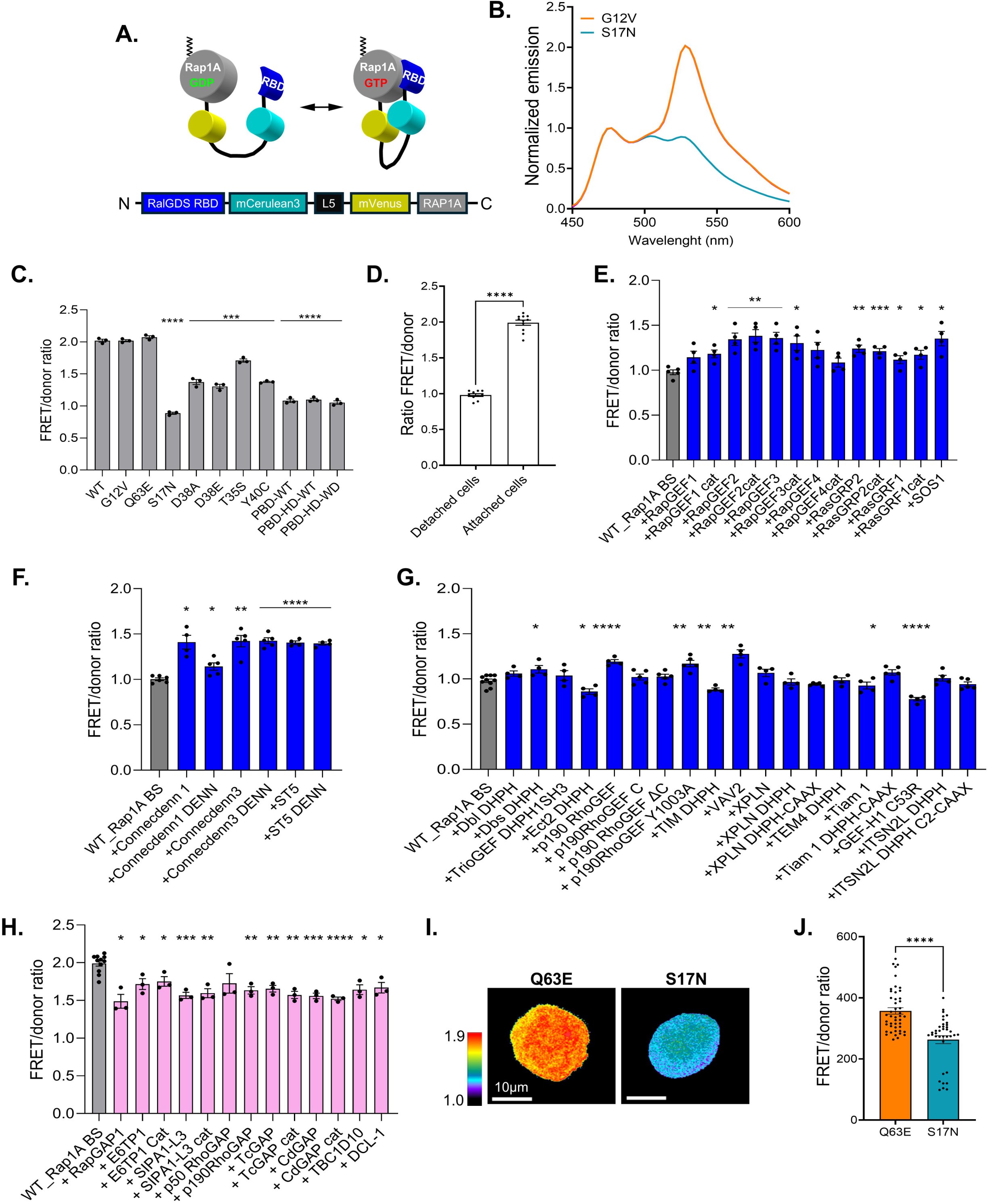
Development and validation of a genetically encoded Rap1A FRET biosensor. (A) Schematic representation of the Rap1A single-chain FRET biosensor. The biosensor consists of fluorescent proteins mCerulean3 and mVenus, flanked by C-terminally attached, full length Rap1A and the N-terminal Ras association domain (RBD) of RalGDS. (B) Representative emission spectra following donor excitation in cells expressing constitutively active (G12V) or inactive (S17N) RAP1A mutant biosensor. (C) Quantification of Rap1A biosensor activity measured in constructs carrying mutations affecting Rap1A activation or the effector-binding interface, n=3. (D) Comparison of Rap1A biosensor activity in adherent and suspension conditions. n=10-11. (E) Rap1A biosensor activity measured following expression of Ras- and Rap-family guanine nucleotide exchange factors (GEFs), n=4-5. (F) Rap1A biosensor activity measured following expression of Rab-family GEFs, n=4. (G) Rap1A biosensor activity measured following expression of Rho-family GEFs, n=3-9. (H) Rap1A biosensor activity measured following expression of Rap GTPase-activating proteins (GAPs) n=3-11. (I) Representative pseudocolor images of mutant Rap1A biosensor activities in cells expressing constitutively active (Q63E) or inactive (S17N) versions of the biosensor. Warmer colors correspond to higher FRET ratios. Scale bar, 10 µm. (J) Quantification of (I). Data represent mean ± SEM, n=3. Each dot represents a single cell. Statistical significance was determined using an unpaired two-tailed Student’s t-test. *p<0.05, **p<0.01, ***p<0.001, ****p<0.0001

Because Rap1 is a well-established regulator of integrin-mediated adhesion (7, 39, 45, 56–58), we next examined whether Rap1A biosensor activity depends on the adhesive state of cells. Fluorometric measurements performed in LinXE cells transiently expressing the Rap1A biosensor revealed that adherent cells displayed approximately twofold higher Rap1A activity than cells maintained in suspension (Fig. 2D). We next examined the response of the biosensor to regulatory proteins that control small GTPase activity. To maintain an optimal dynamic range of biosensor activation, these experiments were performed in detached LinXE. As expected, expression of Rap-specific guanine nucleotide exchange factors (GEFs) strongly increased Rap1A biosensor activity, indicating that the biosensor responds to regulators of endogenous Rap1A (Fig. 2E). Interestingly, Ras-specific GEFs also increased Rap1A biosensor activity (Fig. 2E), consistent with the strong sequence and structural homology between Ras and Rap GTPases, which share several regulators and effectors (52). We next examined the effects of GEFs associated with other small GTPase families. Expression of Rab GEFs involved in endosomal trafficking increased Rap1A biosensor activity (Fig. 2F). Similarly, expression of Rho-family GEFs altered Rap1A biosensor activity, resulting in both activation and inhibition of the biosensor (Fig. 2G). Consistent with these observations, expression of GTPase-activating proteins (GAPs) targeting Rap, Ras, Rab, or Rho GTPases generally reduced Rap1A biosensor activity (Fig. 2H). We next examined differences in biosensor activity between constitutively active and inactive Rap1A mutants using high-resolution microscopy in the rat adenocarcinoma cell line MTLn3. Cells were transiently transfected with Rap1A biosensors harboring either the constitutively active Q63E mutation or the inactive S17N mutation. Biosensor activity was quantified by wide-field fluorescence microscopy using ratiometric analysis (Fig. 1I). We observed a significant reduction in the FRET/donor ratio in the S17N mutant compared with the constitutively active Q63E mutant, confirming the robust dynamic range of the biosensor (Fig. 1J). Together, these results demonstrate that the Rap1A biosensor reports Rap1A activation in living cells and responds to both physiological stimuli and regulatory inputs from multiple small GTPase pathways.

### ARHGAP20 regulates the coordination of Rap1A and RhoA activities in adhesion-associated regions behind the leading edge

To determine how ARHGAP20 regulates the coordination between Rap1A and RhoA during cell migration, we analyzed the spatiotemporal dynamics of both GTPases at the leading edge using live-cell imaging with genetically encoded biosensors. To enable simultaneous monitoring of both signaling activities, we generated a stable MTLn3 cell line co-expressing the Rap1A FRET biosensor and a near-infrared (NIR) RhoA FRET biosensor (Supplemental Fig. S3) derived from a previously published CFP/YFP-based RhoA FRET biosensor (59). When both biosensors were expressed in a single cell, they revealed spatially distinct and heterogeneous activity patterns (Fig. 3A; Supplementary Movie1). To ensure that biosensor expression did not perturb endogenous signaling, we compared the levels of biosensor expression with endogenous proteins. Western blot analysis showed that both Rap1A and RhoA biosensors were expressed at levels substantially lower than their endogenous counterparts (Fig. 3B). Quantification confirmed that induction of biosensor expression did not alter the expression of endogenous Rap1A or RhoA (Fig. 3C), indicating that the biosensors were expressed under conditions compatible with reliable signaling measurements (59, 60). To quantify GTPase activity relative to cell edge dynamics, we used the previously described approach of sampling the Rap1A and RhoA activities in a dynamic coordinate system of probing windows that follow the cell edge movement (61) (Fig. 3D, E). This allowed us to quantify by cross-correlation analysis the temporal relationship between Rap1A and RhoA signaling in different layers of probing windows. Under control conditions, Rap1A and RhoA activities displayed a predominantly negative cross-correlation with strong negative time lags, indicating that activation of the two GTPases occurs in opposite phases and with strong temporal shifts during cycles of edge protrusion and retraction (Fig. 3F). Interestingly, depletion of ARHGAP20 enhanced this negative correlation without strongly affecting the temporal coupling, specifically within window layers located 0.9 to 2.7µm from the cell edge. In control cells, the cross-correlation amplitude remained below the significance threshold (∼0.14), whereas ARHGAP20 depletion increased the cross-correlation peak above this threshold (Fig. 3F). Quantification of cross-correlation peak amplitudes confirmed an increase in Rap1A–RhoA anticorrelation upon ARHGAP20 depletion in these regions (Fig. 3G). Together, these results indicate that ARHGAP20 depletion enhances the negative coordination between Rap1A and RhoA activities in regions that correspond to adhesion-associated zones (62, 63).

**Fig. 3.**
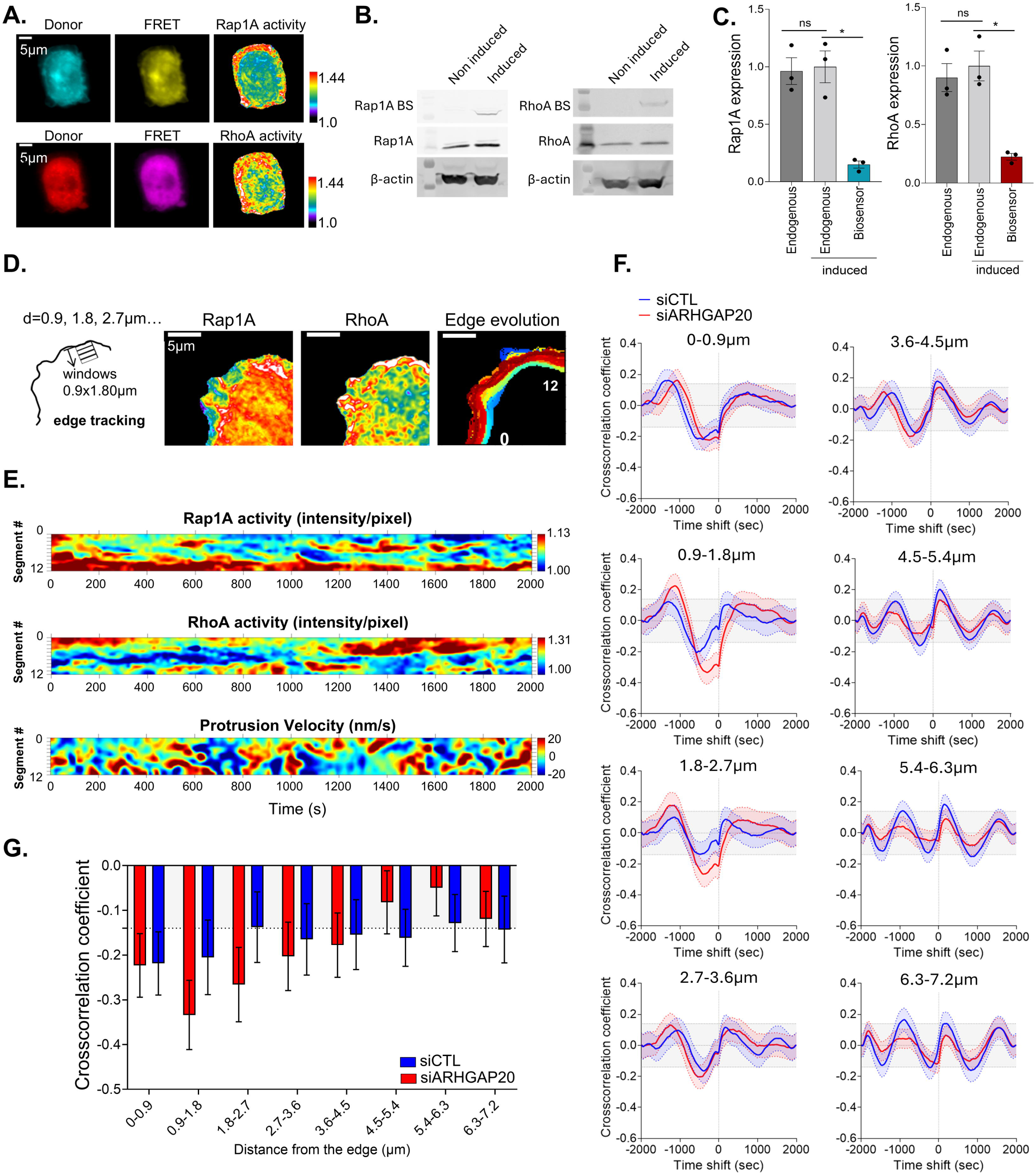
ARHGAP20 regulates the coordination of Rap1A and RhoA activities in adhesion-associated regions behind the leading edge. (A) Representative images of Rap1A and RhoA biosensor activities in motile MTLn3 cells. Cells stably co-expressing the Rap1A FRET biosensor and a near-infrared (NIR) RhoA FRET biosensor were imaged during migration. For Rap1A, cyan (donor), yellow (acceptor-FRET), and ratiometric images are shown. For RhoA, red (donor), magenta (acceptor-FRET), and ratiometric images are shown. Ratiometric pseudo color bars are also shown. Scale bar, 5 µm. (B) Western blot analysis showing expression levels of Rap1A and RhoA biosensors compared with endogenous proteins in MTLn3 cells. Uncropped Westernblots are shown in Supplementary Figure 6. (C) Quantification of (B), n=3. Statistical significance was determined using unpaired two-tailed Student’s *t*-tests. *p<0.05. (D) Schematic representation of the window-based analysis used to quantify biosensor activity relative to the cell edge. The leading edge was sampled in 12 windows following the outline of the plasma membrane (illustrated in the Edge evolution image); each window was divided into 8 segments according to their distance from the cell edge, the first originating 4 pixels behind the cell edge (0.9-1.8um) and the last extending 32 pixels into the cell interior (6.3-7.2 um). Representative ratiometric maps of Rap1A and RhoA activities at the cell edge, and the cell edge tracking evolution are shown. Scale bar, 5 µm (E) Representative heat maps showing the spatiotemporal dynamics of Rap1A activity (top), RhoA activity (middle), and the protrusion velocity (bottom) along the cell edge. The horizontal axis represents time, whereas the vertical axis corresponds to spatial position along the segmented cell edge windows. Color intensity indicates the relative level of biosensor activity or protrusion velocity according to the indicated color scale. Warm colors represent higher activity or protrusion, whereas cool colors represent lower activity or retraction. (F) Cross-correlation analysis between Rap1A and RhoA biosensor activities calculated for each edge window as a function of timelags in siCTL and siARHGAP20 cells. The gray shaded region indicates cross-correlation values below the significance threshold (±0.14). (G) Quantification of cross-correlation peak amplitudes for Rap1A and RhoA activities across the different edge window segments. Data represent mean ± 95%CI. siCTL= n=4, 16 cells, 489 windows, siARHGAP20= n=3, 17 cells, 477 windows.

### ARHGAP20 regulates cell adhesion and Rap1A–RhoA coordination outside focal adhesions

In view of a spatially specific function of ARHGAP20 in zones of focal adhesions, we next examined whether ARHGAP20 depletion alters cell adhesion. Indeed, depletion of ARHGAP20 significantly increased cell adhesion (Fig. 4A). To investigate how Rap1A and RhoA activities are organized relative to adhesion structures, we visualized biosensor activities together with vinculin as the focal adhesion marker (Fig. 4B). We then quantified the spatial relationship between Rap1A and RhoA activities using Pearson’s correlation analysis. At the whole-cell level, no significant difference in Rap1A–RhoA correlation was observed between control and ARHGAP20-depleted cells (Fig. 4C). Similarly, when the analysis was restricted specifically to focal adhesion regions, no significant change in Rap1A–RhoA correlation was detected (Fig. 4D). Because Pearson’s analysis does not allow us to evaluate the degree to which ARHGAP20 may influence the spatiotemporal coordination of both Rap1A and RhoA signaling dynamics and heterogeneity, next we used a recently developed computational method to identify signaling microdomains (MDs) (53), which are regions of the cell with elevated and spatially constrained GTPase signaling coordination. Briefly, the computational pipeline integrates measurements of local signaling coordination as measured by dynamic time warping (DTW) distance to create a ‘coordination movie’ from which spatiotemporal MDs can be extracted (Fig. 4E,F) (53). Unlike standard correlation, DTW is a versatile metric for the similarity of two time series even if they are not aligned or moving at differing speeds (64, 65). We computed MDs separately in Rap1A and RhoA channels for each of the conditions (siCTL and siARHGAP20) and analyzed MD strength, duration, abundance, and cell area coverage. We found strong differences in the strengths of coordination inside the MDs of the two signals (Fig.4G). Regardless of the expression of ARHGAP20, RhoA is more than 1.5x more coordinated than Rap1A. This may be related to more variable regulatory inputs upstream of Rap1A than upstream of RhoA. Other metrics per MD were not statistically different between siCTL and siARHGAP20 (Fig.S5). We then turned to the spatiotemporal colocalization of Rap1A and RhoA MDs, as quantified by the Jaccard Index (area overlap over union of MDs identified for the two signals). When the analysis was performed across the entire cell, ARHGAP20 depletion resulted in a significant reduction of Rap1A–RhoA MD spatiotemporal colocalization compared to the control (Fig. 4H; Supplementary Movies 2 and 3). When we parsed this data for MDs that are associated with focal adhesions as labelled by fluorescent vinculin, the colocalization between Rap1A and RhoA MDs was not significantly affected by ARHGAP20 depletion (Fig. 4I), in contrast to the colocalization in regions excluding the focal adhesions (Fig. 4J). Together, these results indicate that ARHGAP20 depletion decreases the overlap between Rap1A and RhoA signaling outside focal adhesion regions suggesting a misalignment in the antagonism of these two signals.

**Figure 4.**
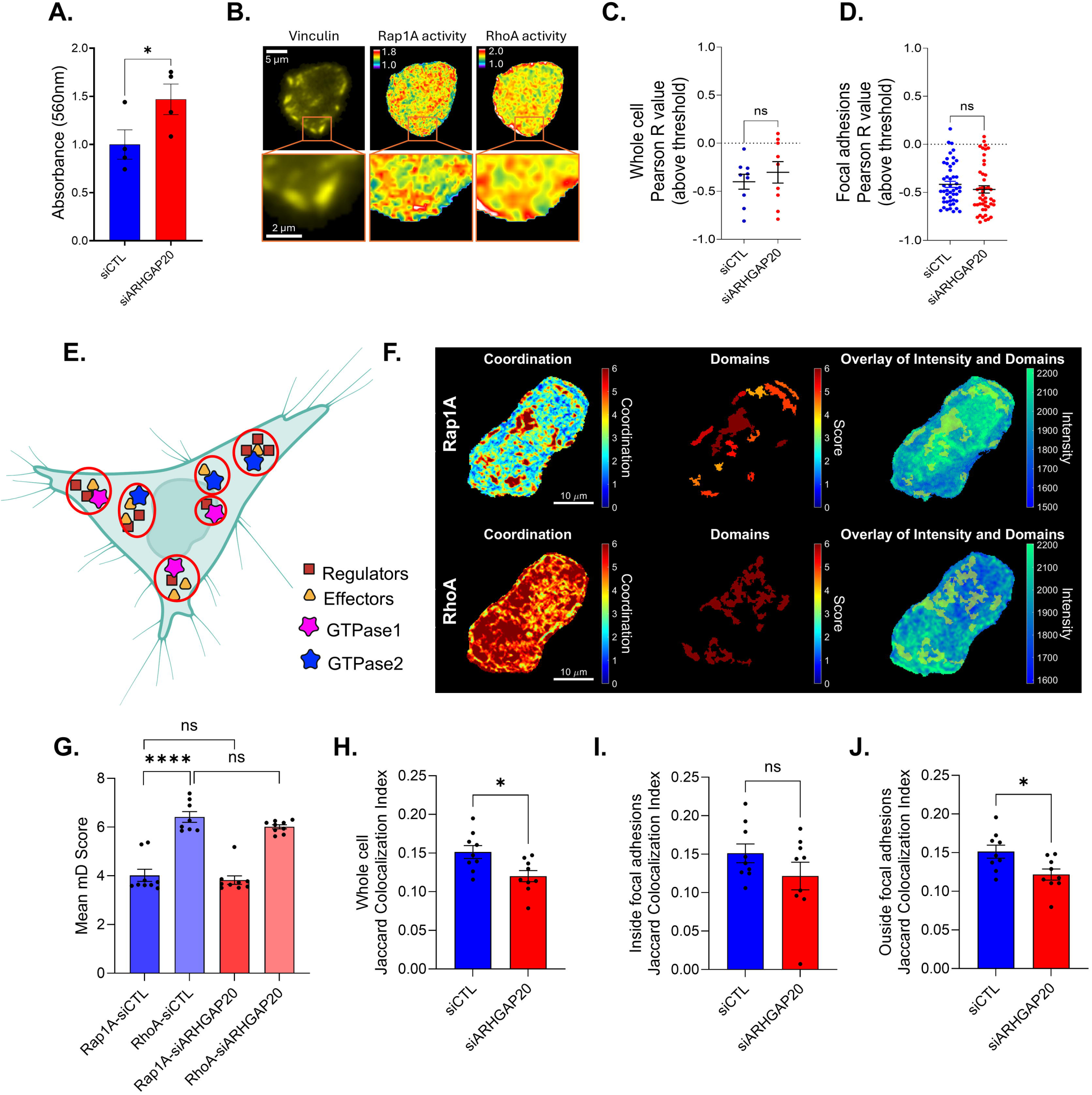
Spatial analysis of Rap1A and RhoA activities relative to focal adhesions. (A) Quantification of adherent MTLn3 cells transduced with either siCTL or siARHGAP20. Cells were allowed to adhere to fibronectin-coated plates, and adhesion was measured by crystal violet staining. Absorbance at 560 nm represents the amount of adherent cells. Data are presented as mean ± SEM from n = 4 independent experiments. (B) Representative images showing Rap1A and RhoA biosensor activity in cells at the focal adhesion. Mtln3 cells expressing the Rap1A and RhoA biosensors were transfected with mRuby3-Vinculin to visualize focal adhesions. (C) Pearson correlation analysis between Rap1A and RhoA biosensor activities calculated across the entire cell. (D) Pearson correlation analysis between Rap1A and RhoA biosensor activities restricted to focal adhesion regions. (E) A cartoon depiction of the microdomains (MDs). A microdomain is conceptualized as a region in the cell with spatially constrained, highly coordinated Rho GTPase signaling activity that is driven by the localization of a set of GEF, GAPs, and effectors. Created with BioRender.com. (F) Representative timepoint images of the computed Coordination, Domains (MD), and the overlay onto the biosensor activity Intensity map, for Rap1A (TOP) and RhoA (BOTTOM) from the siCTL condition. (G) Mean MD Score, representing the strengths of the DTW coupling during MD evolution. (H) Time averaged Jaccard Colocalization index calculated from the MD analysis between Rap1A and RhoA activities calculated across the entire cell. (I) Time averaged Jaccard colocalization index calculated from the MD analysis between Rap1A and RhoA activities within the mRuby3-Vinculin focal adhesions. (J) Time averaged Jaccard colocalization index calculated from the MD analysis between Rap1A and RhoA activities in regions outside mRuby3-Vinculin focal adhesions. Data represent mean ± SEM. Statistical significance was determined using unpaired two-tailed Student’s t-tests. *p<0.05, ****p<0.001, pooled datasets from n=3 independent experiments.

### ARHGAP20 subcellular localization

To determine how ARHGAP20 regulates Rap1A–RhoA coordination outside focal adhesions, we next investigated its subcellular localization by immunofluorescence microscopy using fluorescent markers of distinct intracellular compartments (Fig. 5). Colocalization was quantified using Pearson’s correlation coefficient, and a Pearson’s R value greater than 0.5 was considered indicative of strong colocalization. We first examined the localization of ARHGAP20 relative to the focal adhesion marker vinculin (Fig. 5A, B). Consistent with our microdomain analysis, ARHGAP20 displayed no colocalization with vinculin, indicating that it is not enriched within focal adhesions themselves despite regulating Rap1A–RhoA coordination in their vicinity. Because ARHGAP20 acts outside focal adhesions, we next investigated its association with the cytoskeleton. ARHGAP20 exhibited minimal colocalization with filamentous actin but showed significant colocalization with microtubules (Fig. 5C, D), suggesting that ARHGAP20 preferentially localizes within the inner region of the cell rather than at the actin-rich cell periphery. We next assessed the localization of ARHGAP20 relative to intracellular organelles. ARHGAP20 exhibited strong colocalization with both the endoplasmic reticulum and the Golgi apparatus but not with the mitochondria (Fig. 5E–G), supporting its association with the secretory pathway. To further characterize this localization, we analyzed markers of vesicular trafficking. ARHGAP20 strongly colocalized with Rab11- and Rab14-positive recycling and exocytic vesicles, as well as with Rab5-and Rab7-positive endocytic compartments, whereas little colocalization was observed with the exocytic GTPase TC10 (Fig. 5G, H). Importantly, ARHGAP20 lacks an obvious hydrophobic transmembrane domain, making direct membrane insertion unlikely, while its Plekstrin-homology domain may selectively enable interaction with specific phosphoinositol lipids (43, 66). Together, these observations support a model in which ARHGAP20 is recruited to the cytosolic surface of trafficking vesicles. Such localization would allow ARHGAP20 to reach specific regions of the plasma membrane through vesicular trafficking, where it could locally regulate RhoA activity before being recycled through the endocytic pathway, or alternatively, by analogy with the endocytic trafficking described for Rac1 (67, 68), activated RhoA may undergo endocytic uptake and encounter vesicle-associated ARHGAP20 within the recycling compartment, where ARHGAP20 could promote local RhoA inactivation prior to its extraction and sequestration by RhoGDI. The working model is summarized in Fig. 5I.

**Figure 5.**
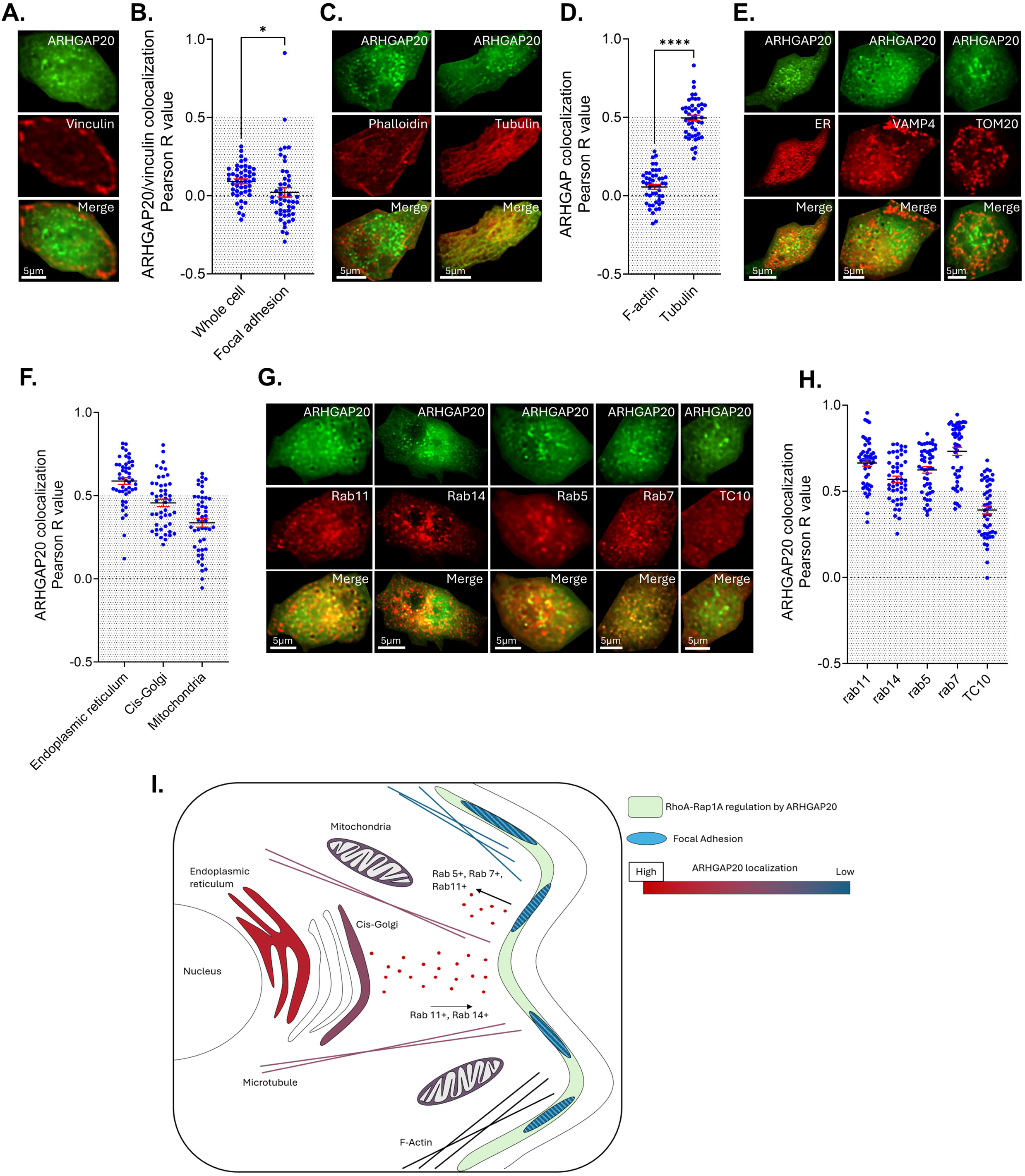
ARHGAP20 subcellular localization. (A) Representative immunofluorescence images showing ARHGAP20 (top), vinculin (middle), and the corresponding merged image (bottom) in MTLn3 cells. (B) Pearson’s correlation analysis of the colocalization between ARHGAP20 and vinculin calculated across the entire cell or restricted to focal adhesion regions. (C) Representative immunofluorescence images showing ARHGAP20 (top), F-actin (left, phalloidin staining) or α-tubulin (right) (middle), and the corresponding merged images (bottom). (D) Pearson’s correlation analysis of the colocalization between ARHGAP20 and F-actin or α-tubulin calculated across the entire cell. (E) Representative immunofluorescence images showing ARHGAP20 (top), the endoplasmic reticulum (ER), VAMP4, or TOM20 (middle), and the corresponding merged images (bottom), from left to right. (F) Pearson’s correlation analysis of the colocalization between ARHGAP20 and the ER, VAMP4, or TOM20 calculated across the entire cell. (G) Representative immunofluorescence images showing ARHGAP20 (top), Rab11, Rab14, Rab5, Rab7, or TC10 (middle), and the corresponding merged images (bottom), from left to right. (H) Pearson’s correlation analysis of the colocalization between ARHGAP20 and Rab11, Rab14, Rab5, Rab7, or TC10 calculated across the entire cell. Data are presented as mean ± SEM. Statistical significance was determined using unpaired two-tailed Student’s t-tests *p<0.05, ****p<0.0001. Pooled datasets from n = 3 independent experiments, with 15–16 cells analyzed per experiment. (I) Proposed model summarizing the role of ARHGAP20 in the spatial organization of Rap1A–RhoA signaling and the regulation of focal adhesion dynamics and directional cell migration.

## Discussion

Cell migration requires the precise spatial and temporal coordination of signaling pathways that regulate cytoskeletal dynamics and adhesion turnover. In this study, we identify ARHGAP20 as a regulator of cell migration that modulates the coordination between Rap1A and RhoA signaling. Using quantitative migration assays and live-cell imaging of multiplex, genetically encoded biosensors, we show that depletion of ARHGAP20 reduces migratory capacity, increases cell adhesion, and alters the spatial coordination between Rap1A and RhoA activities in regions outside focal adhesions.

Consistent with previous reports linking ARHGAP20 to cancer cell motility (41), partial depletion of ARHGAP20 significantly impaired both directed and random migration of MTLn3 cells. ARHGAP20-depleted cells displayed reduced velocity and displacement, remaining confined near their point of origin during long-term tracking. These defects were accompanied by increased cell adhesion, suggesting that ARHGAP20 contributes to the regulation of adhesion dynamics required for efficient migration. Because productive migration requires a balance between adhesion formation and turnover (9, 69–74), excessive adhesion can restrict cell movement by limiting adhesion detachment or by stabilizing adhesion structures that normally undergo rapid remodeling during migration.

To investigate the signaling mechanisms underlying these phenotypes, we developed and validated a genetically encoded, single-chain FRET biosensor for Rap1A. This biosensor enabled direct visualization of Rap1A activation in living cells and responded robustly to multiple classes of regulators, including Rap-, Ras-, Rab-, and Rho-family GEFs and GAPs. These observations highlight the integration of Rap1A signaling within broader small GTPase networks that regulate membrane trafficking, cytoskeletal dynamics, and adhesion signaling (7, 8, 20, 36, 47, 54, 75–77). The ability of the biosensor to detect adhesion-dependent Rap1A activation further supports the well-established role of Rap1A signaling in integrin-mediated adhesion and adhesion maturation (7, 39, 78).

Using simultaneous imaging of Rap1A and RhoA activities, we observed that the two GTPases exhibit predominantly negative temporal coordination during protrusion cycles at the cell edge. Such negative coordination is consistent with the distinct functions of these signaling pathways during migration, where Rap1A activity promotes integrin activation and adhesion formation (7, 39, 45, 46), while RhoA regulates actomyosin contractility and adhesion maturation (9, 14, 21, 36, 47, 48). The temporal separation of their activities may therefore contribute to the cyclical regulation of protrusion, adhesion assembly, and contractile forces that drive cell movement.

Importantly, depletion of ARHGAP20 significantly enhanced the negative coordination between Rap1A and RhoA signaling in regions located just behind the leading edge. These regions correspond to areas enriched in focal adhesion structures rather than the extreme protrusive edge. Interestingly, this effect of ARHGAP20 depletion on negative cross-correlation was not detected within focal adhesions themselves. However, MD analysis revealed that, in regions outside focal adhesions, ARHGAP20 depletion reduced the degree of spatiotemporal overlap between Rap1A and RhoA signaling complexes. These findings suggest that ARHGAP20 does not act within adhesion structures *per se* but instead regulates signaling organization and dynamics in adjacent regions that influence adhesion behavior. The spatial restriction of ARHGAP20-dependent signaling effects outside focal adhesions suggests that ARHGAP20 may control how signaling gradients propagate between protrusive and adhesive regions of migrating cells. By modulating the coordination between Rap1A and RhoA activities and MDs in these zones, ARHGAP20 may contribute to the fine-tuning of protrusive dynamics and adhesion assembly and turnover required for efficient migration. In this context, increased anticorrelation between Rap1A and RhoA activities upon ARHGAP20 depletion may reflect an imbalance in signaling dynamics that stabilize adhesions and restrict cell motility. Our study identifies vesicular trafficking as a previously unrecognized component of ARHGAP20-mediated regulation of RhoA signaling. Although ARHGAP20 regulates Rap1A–RhoA coordination in microdomains surrounding focal adhesions, it is itself largely absent from focal adhesions, indicating that its activity is spatially controlled before reaching these signaling sites. Instead, ARHGAP20 preferentially associates with microtubules and intracellular membrane compartments, including the endoplasmic reticulum, the Golgi apparatus, and multiple Rab-positive vesicular populations. The strong colocalization with Rab11 and Rab14 suggests that ARHGAP20 is transported through the secretory and recycling pathways, whereas its association with Rab5 and Rab7 indicates subsequent entry into the endocytic pathway. In contrast, the absence of significant colocalization with TC10 argues against a generalized association with all exocytic vesicles and instead suggests selective recruitment to specific trafficking routes. Because ARHGAP20 does not contain an obvious transmembrane domain, our data support a model in which ARHGAP20 decorates the cytosolic surface of trafficking vesicles, likely through association with phosphoinositides via its PH domain (43, 66). This organization provides an attractive mechanism to spatially restrict GAP activity by delivering ARHGAP20 close to the plasma membrane, where RhoA is plasma membrane-associated and active. Following local RhoA inactivation, either through direct interaction with ARHGAP20 at the plasma membrane or, hypothetically, following endocytic uptake of RhoA into the recycling pathway where it may encounter ARHGAP20, the GAP itself could subsequently be removed from the cell cortex through endocytic trafficking, thereby restricting the spatial and temporal extent of RhoA inhibition. Such a trafficking-dependent mechanism would explain how ARHGAP20 regulates Rap1A–RhoA coordination adjacent to focal adhesions without being stably associated with the adhesions themselves. More broadly, these findings support the emerging concept that intracellular trafficking not only transports signaling molecules but also establishes the spatial organization of signaling networks controlling cell migration.

Taken together, our results support a model in which ARHGAP20 regulates cell migration by modulating the spatial coordination of Rap1A and RhoA signaling in regions surrounding focal adhesions. Through this mechanism, ARHGAP20 may help control the balance between adhesion stabilization and cytoskeletal contractility that underlies efficient cell movement. More broadly, these findings highlight the importance of spatially organized crosstalk between small GTPase signaling networks in controlling the dynamic behaviors required for cell migration. Beyond its role in regulating migration *in vitro*, our findings may also have implications for tumor cell dissemination. ARHGAP20 was previously reported to be upregulated in early disseminating tumor cells, a population characterized by enhanced motility and invasive potential (44). In this context, the ability of ARHGAP20 to modulate the coordination between Rap1A and RhoA signaling may contribute to the dynamic regulation of adhesion and cytoskeletal contractility required for invasive migration. Efficient tumor dissemination requires cells to continuously adjust adhesion strength while maintaining productive protrusion and contractile forces, allowing cells to navigate through heterogeneous extracellular environments (8, 44, 79–83). By tuning the spatial coupling between Rap1A-driven adhesion signaling and RhoA-mediated contractility, ARHGAP20 may facilitate the transitions between adhesive and migratory states that enable invasive behavior. Future studies will therefore be required to determine whether ARHGAP20-dependent regulation of Rap1A–RhoA coordination contributes to tumor cell invasion and metastatic dissemination in more complex physiological environments.

## Material and methods

### Cell culture

MTLn3 cells (84) (rat mammary adenocarcinoma) were cultured in Minimum Essential Medium (MEM, Corning, Corning, NY, USA) supplemented with 5% fetal bovine serum (FBS), 1% glutamine, and 100 I.U. penicillin and 100Oµg/mL streptomycin (Invitrogen, Carlsbad, CA, USA), as previously described (85). LinXe (HEK-293T derivative; gift from Dr.Klaus Hahn (86)) cells were cultured in Dulbecco’s modified Eagle medium (DMEM, Corning, Corning, NY, USA) supplemented with 10% FBS, 1% glutamine, and penicillin/streptomycin, as previously described (87, 88).

### Plasmid transfections

MTLn3 cells were plated at a density of 2 × 10O cells per well in 6-well plates and allowed to adhere overnight under standard culture conditions. The following day, 2 µg of plasmid DNA was diluted in 250 µL of Opti-MEM, while 4 µL of Lipofectamine 2000 (Life Technologies, Carlsbad, CA, USA) was diluted separately in 250 µL of Opti-MEM and incubated for 5 minutes at room temperature. The diluted Lipofectamine was then combined with the DNA solution and incubated for 20 minutes at room temperature to allow complex formation. Cells were washed once with PBS and incubated with 500 µL of Opti-MEM. The transfection mixture was then added to the cells. After 45 minutes, the medium was replaced with complete growth medium, and cells were further incubated overnight prior to downstream analyses.

### Expression constructs and plasmids

Following expression constructs were used: pmRuby3-C3, pmRuby3-C3-Vinculin, miRFP680-C3-Vinculin (all vinculin constructs were based on a gift from Dr. Klaus Hahn), EGFP-alpha-Tubulin (a gift from Dr. Claire Waterman), TOM20-miRFP680, ER-tdTomato (gift from Dr. Erik Snapp), VAMP4-EGFP (a gift from Dr. Thierry Galli; Addgene #42313), Rab11-mCherry (a gift from Dr. Michael Davidson; Addgene # 55124), mNeonGreen-C3-Rab14 (based on a gift from Dr. Curt Civin; Addgene #61493), mNeonGreen-Rab5 (based on a gift from Dr. Michael Davidson; Addgene #55126), mCherry-Rab7(a gift from Dr. Michael Davidson; Addgene #55127), mCherry-TC10 (89), and FRET biosensors. A codon optimized mRuby3 (90) was synthesized (Azenta, Burlington, MA, USA) to produce the pmRuby3-C3 construct. A codon optimized miRFP680 (91) was synthesized (Azenta, Burlington, MA, USA) to produce pmiRFP680-C3 construct. A codon optimized mNeonGreen (92) was synthesized (Azenta, Burlington, MA, USA) to produce the pmNeonGreen-C3 construct. pmRuby3-C3-Vinculin and pmiRFP680-C3-Vinculin was constructed by PCR amplifying chicken Vinculin (93), using the primer pair: 5’-GGAATAATATAGAACAAGCTTATGCCCGTCTTCCACACGCGCACCA-3’ and 5’-GCAAACTTTCATATAGAATTCTTACTGATACCATGGGGTCTTTCTGACCCA-3’, and ligated into backbones at HindIII/EcoRI sites. TOM20-miRFP680 was constructed by two-step PCR using the primer pair to amplify TOM20: 5’-GGATAATTATAATTACCATGGTGGGCCGCAACAGCGCGATTG-3’; and 5’-TGGCGGGCCACGCTGCCCTCGGCCATTCTAGATTTAAAGTTCGGATCGCT-3; and miRFP680: 5’-AGCGATCCGAACTTTAAATCTAGAATGGCCGAGGGCAGCGTGGCCCGCCA-3’ and 5’-GGAATTATAATTAAACTCGAGTTACTCCTCCATCACGCCGATCTGAGTG-3’, and ligated into NcoI/XhoI sites in pTriEX-4 backbone (Sigma Aldrich, St.Louis, MO, USA). mNeonGreen-C3-Rab14 were constructed by PCR amplifying Rab14 using the primer pair: 5’-GGAACTATCTATAGAATCAACTCGAGATGGCAACTGCACCATACAACTACTCT-3’ and 5’-GCATATGGAATAACAAGGATCCCTAGCAGCCACAGCCTTCTCTCTGGGGTTGGGGT-3’, and ligated into BamHI/XhoI sites of the pmNeonGreen-C3 backbone. Rab5A was restriction digested at XhoI/BamHIsites and ligated into pmNeonGreen-C3 backbone.

### FRET biosensors and stable cellular expression by viral transduction

Cyan-yellow fluorescent protein-based FRET biosensor for Rap1A and B were constructed as following: Human RalGDS RBD (786-883 amino acids) was used for the affinity moiety, and PCR amplified using the primer pair: 5’-GCCTAGAACTGTACTTATCA CCATGGCCTCCGCGCTGCCGCTCTACAACCAGC-3’ and 5’-GCATCCGTACAGTTCGAGTCGGATCCGGTCCGCTTCTTCAGGACAAAGTCA-3’, and ligated into int pTriEX-4 at NcoI/BamHI sites. Synonymous codon modified mCerulean3 was PCR amplified using the primer pair: 5’-GCATTGCATAGCATATAGAAGGATCCGGAATGGTGTCCAAAGGAGAAGAACTGT-3’ and 5’-GCTTAATATGCATATATATCAAGCTTTTATACAGTTCATCCATTCCCAGG-3’, and ligated into BamHI/HindIII sites of the pTriEX-RalGDS-RBD backbone. The linker segment of 93 amino acids contained 5x repeating units of a previously published flexible linker design (94), was synonymous codon modified and synthesized (Azenta, Burlington, MA, USA), then ligated into HindIII/NotI sites within the pTriEX-RalGDS-RBD-mCer3 backbone. Monomeric Venus was restriction digested at NotI/EcoRI from a previous Rab13 FRET biosensor (95) and ligated into the pTriEX-RalGDS-RBD-mCer3-5xLinker backbone. Rap1A or B was PCR amplified using the primer sets: 5’-GCAATCTTACGAACTGTTAAGAATTCATGCGTGAGTACAAGCTAGTGGTCC-3’ and 5’-GCTGAAATTGATTCGAATTTCTCGAGCTAGAGCAGCAGACATGATTTCTTTTTAG-3’ for Rap1A and 5’-GGTATAATATTAATAGAATTCATGCGTGAGTATAAGCTAGTCGTTC-3’ and 5’-GGTTTTAATCATAAACTCGAGTCAAAGCAGCTGACATGATGACTTTTT-3’ for Rap1B, and ligated into EcoRI/XhoI sites of the biosensor backbone.

For NIR FRET RhoA biosensor, codon optimized miRFP720 (96) was PCR amplified using a primer pair: 5’-GGATAATTCAAGTTCACCATGGCCGAGGGCAGCGTGGCCCGCCA-3’ and 5’-GCATCATCTTAAGAATGGATCCCTCCTCCATCACGCCGATCTGGGTGG-3’, and ligated into NcoI/BamHI of the pTriEX-4 biosensor backbone. RBD from Rhotekin (59) was 2 step PCR amplified to include a short flexible linker of 6 a. a. followed by a 2x Proline inducing a turn in the structure within the C-terminus of the fragment, by using primer set: 5’-GCATTAAATTATATGCATATGGATCCATCCTGGAGGACCTGAACATGCTGT-3’, 5’-GGGTCCGCTGCCGCCGCTGCCGGTCTTCTCCAGCACCTGGGCCTCC-3’, and 5’-GCAATTACTATTTAGCAATGAAGCTTCCGGGGGGTCCGCTGCCG-3’, and ligated at BamHI/HindIII of the biosensor backbone. A synonymous codon-based, 3x repeating units of flexible linker segment (54 a.a.) (94) was restriction digested from the previous Rab13 biosensor (95) and ligated at HindIII/NotI sites of the biosensor backbone. miRFP670nano was PCR amplified using the primer pair: 5’-GGAATATTTAATGCATATGCGGCCGCAATGGCAAACCTGGACAAGATGCTGA-3’ and 5’-CGTTATATATGCATATTAGAATTCGCTCTGCTGGATGGCGATGCCCACC-3’ and ligated into NotI/EcoRI sites of the biosensor backbone. A full length RhoA was restriction digested from our previous cyan-yellow FRET based biosensor for RhoA (59) and ligated into the EcoRI/XhoI sites of the biosensor backbone.

To construct tetracycline-inducible retroviral expression vectors for the biosensors, the pRetro-X system (Takara Bio USA, San Jose, CA, USA) was adapted for Gateway-based cloning. In brief, pRetro-X-puro and pRetro-X-hygro backbones were engineered by inserting a Gateway destination cassette (Invitrogen) into the multiple cloning site. The Rap1A biosensor and NIR RhoA biosensor cDNA were excised from the pTriEX-4 plasmid by NcoI/XhoI digestion and subcloned into pENTR-4 (Life Technologies, Carlsbad, CA, USA). The resulting entry clones were then recombined with the modified pRetro-X destination vectors using LR Clonase II (Life Technologies, Carlsbad, CA, USA), following the manufacturer’s protocol.

Stable MTLn3 cell lines expressing the biosensors were established by retroviral transduction following standard protocols (97). Rap1A biosensor expression was selected with puromycin (10 ug/mL; Corning, Corning, NY, USA), whereas NIR RhoA biosensor expression was selected with hygromycin B (200 mg/mL; Santa Cruz Biotechnology, Dallas, TX, USA). After antibiotic selection, stable populations expressing both biosensors were further refined by fluorescence-activated cell sorting (FACS) to recover cells with low-to-intermediate biosensor expression levels, thereby limiting artifacts associated with overexpression while preserving adequate fluorescence intensity for live-cell imaging.

Biosensor base-pair sequences are shown in Supplementary Data 1-3.

### siRNA transfection

MTLn3 cells were transfected with siRNA using Oligofectamine (Life Technologies, Carlsbad, CA, USA) according to the manufacturer’s instructions. ARHGAP20 knockdown was achieved using siARHGAP20 (Smartpool, Cat# L-102680-02-005; Dharmacon, Lafayette, CO, USA), and a non-targeting siRNA was used as control (siCTRL; Cat# SIC002; Millipore Sigma, Burlington, MA, USA).

### RTqPCR

Total RNA was extracted from cultured cells using RNeasy plus mini Kit (Cat# 74134; Qiagen, Germantown, MD, USA) according to the manufacturer’s instructions. Total RNA was reverse transcribed into cDNA using High capacity cDNA reverse transcription Kit (Cat# 4368814; Applied Biosystems, Woburn, MA, USA) according to the manufacturer’s protocol. Quantitative PCR was performed using Power up SYBR Green master mix (Cat# A25741; Applied Biosystems, Woburn, MA, USA) on a QuantStudio3 (Applied Biosystems, Woburn, MA, USA). Reactions were carried out in duplicate using gene-specific primers. Relative mRNA expression levels were determined using the 2^−ΔΔCt method (98) and normalized to GAPDH, YWHAZ and PPIA expression. Melt curve analysis was performed at the end of each run to confirm amplification specificity.

Primers :

rGapdh FD 5’ TGCACCACCAACTGCTTAG 3’

rGapDH RV 5’GGATGCAGGGATGATGTTC3

rYwhaz FD 5’GATGAAGCCATTGCTGAACTTG3’

rYwhaz RV 5’GTCTCCTTGGGTATCCGATGTC3’

rPpia FD 5’AGGATTCATGTGCCAGGGTG3’

rPpia RV 5’CTCAGTCTTGGCAGTGCAGA3’

Arhgap20 FD 5’GCCATGAAATCCTTTGTTTTGGGCT3’

Arhgap20 RV 5’GCATAAGTCTTTGCAAATGGAGGGG3’

### Boyden chamber migration assay

Cell migration was assessed using Boyden chamber inserts (Cat# 353097; Corning, Corning, NY, USA). Cells were seeded in the upper chamber in medium containing 0.5% serum, while the lower chamber was filled with medium supplemented with 5% serum to establish a chemotactic gradient. After incubation for the indicated time, non-invading cells were removed from the upper side of the membrane. Invaded cells on the lower surface were stained with NucBlue (Life Technologies, Carlsbad, CA, USA) according to the manufacturer’s instructions and imaged by fluorescence microscopy. The number of invading cells was then quantified using Image J.

### Random migration assay

Random cell migration was analyzed by time-lapse imaging. Cells were plated onto gelatin-coated glass coverslips and allowed to adhere prior to imaging. Live-cell imaging was performed for 6 h using differential interference contrast (DIC) microscopy under standard culture conditions. Cell trajectories were tracked using MetaMorph software (Molecular Devices, Downingtown, PA, USA). Quantitative parameters, including migration distance and cell velocity, were extracted from the tracked trajectories.

### Attachment assay

The attachment assay was performed based on the protocol described by Feoktistova et al. (99). 2.5x10^5^ MTLn3 cells were seeded onto fibronectin-coated 24-wells plates that had been pre-blocked with BSA to prevent non-specific adhesion. Cells were allowed to adhere for 1 h 30 min at 37°C. Non-adherent cells were removed by gentle washing with PBS, and adherent cells were fixed with 3.7% paraformaldehyde. Cells were then stained with crystal violet. After staining, crystal violet was solubilized using acetic acid, and absorbance was measured at 560 nm using a plate reader to quantify cell adhesion.

### Microscopy imaging

MTLn3 cells were seeded onto gelatin-coated glass coverslips at a density of 1.5 × 10O cells per coverslip. For live imaging, cells were maintained in phenol red-free Ham’s F12K medium (Crystalgen, Commack, NY, USA) supplemented with 1× GlutaMAX (Life Technologies, Carlsbad, CA, USA). Before imaging, the medium was aerated with argon gas for approximately 1 min to lower dissolved oxygen. To further limit photobleaching and phototoxicity, the medium was supplemented with 5% fetal bovine serum, Oxyfluor reagent (1:100; Oxyrase Inc., Mansfield, OH, USA), and 10 mM DL-lactate (Sigma Aldrich, St.Louis, MO, USA). Live-cell imaging was performed in a temperature-controlled chamber maintained at 37 °C.

Live-cell fluorescence imaging was performed on a custom multimodal microscope built around an Olympus IX83 platform equipped with the ZDC2 focus-drift compensation system (Evident Scientific, Needham, MA, USA) and a 60× Olympus UIS2 objective (NA 1.5). For multiplex FRET imaging, the system was configured for simultaneous acquisition of donor and FRET emission channels using two Prime BSI Express cooled sCMOS cameras (Photometrics, Tuscon, AZ, USA) coupled through a beam-splitting device for cyan-yellow system, and two Prime BSI cooled sCMOS detectors (Photometrics, Tuscon, AZ, USA) for the NIR system, allowing concurrent acquisition of four emission channels as described previously (100). Excitation was provided by a Sola-UVN solid-state white-light source (Lumencor, Beaverton, OR, USA), with wavelength selection controlled by a motorized filter wheel (Ludl Electronic Products, Hawthorne, NY, USA). ECFP was excited using an ET436/20X filter (Chroma Technology, Bellows Falls, VT, USA), whereas miRFP670nano was excited with FF01-628/32 (AVR Optics, Fairport, NY, USA). Emission was collected through ET480/40M for ECFP, ET535/30M for cyan to yellow FRET, FF01-681/24 for miRFP670, and FF01-794/160 for the NIR FRET. Depending on the imaging configuration, the primary dichroic in the microscope turret consisted of one of the following Chroma filters: ZT440/488/561/635rpc-UF3. Within the emission-splitting path, a T560LPXRXT-UF2 dichroic separated cyan/yellow fluorescence from red and near-infrared signals, and a T505LPXR-UF2 dichroic further separated cyan donor emission from yellow FRET emission. miRFP670nano1 and miRFP720 emissions were separated using a T700LPXR-UF2 dichroic mirror.

Total internal reflection fluorescence (TIRF) imaging was performed using a Visitron Orbital 200 unit integrated with a VS-LMS laser merge system containing 405, 445, 473, 561, and 640 nm solid-state lasers (Visitron Systems, Puchheim, Germany). This module was incorporated into the multichannel microscope, and the appropriate turret dichroic was selected according to the excitation wavelength. For each laser line, the critical angle was calibrated at the back focal plane of the 60×/1.5 NA objective, and the evanescent-field penetration depth was set to 320 nm.

Microscope hardware, stage movement, and image acquisition, including synchronized four-camera operation, were controlled with VisiView version 7.0.0.9 (Visitron Systems, Puchheim, Germany). Camera synchronization was achieved with an external Virtex trigger device interfaced with VisiView. Simultaneously acquired donor and FRET image pairs were registered using *a priori* calibration and affine transformation-based morphing to permit pixel-resolved alignment before ratiometric analysis, as described previously (101). Differential interference contrast (DIC) transmitted-light images were also acquired during live-cell experiments.

### Morphodynamics analysis

Cell edge motion was quantified using the Morphodynamics framework as described previously (61). In brief, analysis was performed on cropped time-lapse sequences containing regions of the cell perimeter that exhibited sustained protrusion–retraction dynamics. Edge tracking was carried out with the protrusion analysis software (61), which generated a series of sampling windows along the leading edge. Each window measured 4 × 8 pixels, corresponding to 0.86 × 1.7 µm at 60× magnification; this size was previously selected to approximate the diffusion-limited sampling area for acquisitions spaced 10 s apart. Depending on the length of the analyzed leading-edge segment, each cell typically yielded 30–100 such windows. To examine spatial coupling relative to the edge, the sampling windows were propagated inward from the cell boundary in 4-pixel increments, allowing correlation analysis to be extended to distances of up to approximately 7 µm from the leading edge. For each window, normalized cross-correlation functions were calculated in MATLAB using *xcov*, comparing directly cross-correlation function between Rap1A and RhoA activity signals. The cross-correlation function obtained for each window was treated as an independent observation, fitted with a smoothing spline, and then pooled across all analyzed cells. The mean time lag corresponding to the maximal cross-correlation coefficient, together with the 95% confidence interval, was estimated using a nonparametric bootstrap approach (102).

### Signaling microdomain analysis

Signaling microdomains (MDs) were identified and tracked using the previously introduced microdomain analysis pipeline (53). For each intensity movie, a corresponding coordination movie was computed by integrating the local time series coordination at each pixel with its neighbors over a predefined set of downsampling levels and bandwidth lengths. The coordination movie provides a moving representation of local time series coordination. Areas with elevated, spatially constrained signaling represent microdomains as in Fig. 4E. To isolate these regions, the corresponding coordination movie was segmented to generate candidate domains. Low-scoring candidate domains were filtered, and remaining candidate domains were tracked over time to generate a final set of spatio-temporal microdomains. The full details of the method can be found in (53).

Analyses were performed over characteristic MD durations of 51 and 99 frames, with image acquisition at 10 s per frame. A minimum microdomain size threshold of 40 contiguous pixels was applied, with no upper size cutoff, and pixel connectivity in the x-y plane was defined using a 4-way connectivity The binary masks of focal adhesions were created by intensity thresholding the top 10% of the histogram-stretched intensity values of the fluorescent-vinculin channel images corresponding to each time point of the analysis where both Rap1 and RhoA activities were measured. The resulting binary masks were then compared using the Jaccard index, defined as J(A, B)=(A Ö B)/(A Ö B) (103), to quantify the percentage of overlap between microdomains and the focal adhesions.

### Immunofluorescence microscopy and colocalization analysis

Cells were transfected with fluorescent compartment markers using Lipofectamine 2000 according to the manufacturer’s instructions. Twenty-four hours after transfection, cells were fixed with 3.7% paraformaldehyde, permeabilized with 0.3% Triton X-100, and blocked in 2.5% BSA and 2.5% serum. Endogenous ARHGAP20 was detected using a rabbit anti-ARHGAP20 antibody (PA5-104102; Life Technologies, Carlsbad, CA, USA), followed by the appropriate fluorescent secondary antibody. Filamentous actin (F-actin) was visualized using Alexa Fluor 594-conjugated phalloidin. Colocalization between ARHGAP20 and the indicated intracellular markers was quantified using Pearson’s correlation coefficient (Pearson’s *R*) calculated with the Colocalization test plugin in Fiji/ImageJ. For vinculin, Pearson’s *R* was measured either across the entire cell or within manually defined focal adhesion regions. For all other markers, colocalization was quantified over the entire cell using identical analysis parameters across all conditions. Individual cells were analyzed independently, and each data point represents one cell. Three independent biological experiments were performed, with 15–16 cells analyzed per experiment. For graphical representation, a Pearson’s *R* value greater than 0.5 was used as an empirical threshold to indicate strong colocalization. This threshold was used for visualization purposes only and was not included in the statistical analysis. Statistical comparisons were performed using an unpaired two-tailed Student’s *t*-test, and data are presented as mean ± SEM.

## Supplementary figure legend

**Supplemental Fig. S1 : Rap1A biosensor pull down**

Rap1A biosensor GST-RBD-pulldown experiment. LinXE cells were transfected with the following: Lanes 1, untransfected control; 2, mCer1-Rap1A Q63E; 3, mCer1-Rap1A S17N; 4, full length biosensor with Q63E mutant Rap1A; and 5, full length biosensor with Q63E mutant Rap1A and PAK1PBD domain exchanged for the RalGDS-RBD domain.

**Supplemental Fig S2 : Rap1A and RhoA biosensor activities**

Whole cell Rap1A biosensor activity, with siCTL or siARHGAP20. (B) Whole cell RhoA biosensor activity, with siCTL or siARHGAP20. (C) Rap1A biosensor activity at the leading egde, with siCTL (BLUE) or siARHGAP20 (RED). (D) RhoA biosensor activity at the leading, with siCTL (BLUE) or siARHGAP20 (RED). Data represent mean ± SEM, in siCTL (n=7), siARHGAP20 (n=9). Statistical significance was determined using unpaired two-tailed Student’s t-tests.

**Supplemental Fig. S3 : Development and validation of a genetically encoded NIR RhoA FRET biosensor**

Schematic representation of the NIR RhoA single-chain FRET biosensor. The biosensor consists of a C-terminal, full length RhoA and an internally inserted Rho binding domain (RBD) of Rhotekin, with a FRET pair of NIR fluorescent protein and an optimized linker as shown. (B) Representative emission spectra following donor excitation in cells expressing constitutively active (Q63L) or dominant-negative (T19N) RhoA mutant biosensor. (C) Quantification of RhoA biosensor activity measured in biosensor constructs carrying mutations affecting RhoA activation or the effector-binding interface. (D) Comparison of RhoA biosensor activity in adherent and suspension conditions, with or without siARHGAP20, n=4-10. (E) RhoA biosensor activity measured following expression of GTPase-activating proteins (GAPs), n=3-11. (F) RhoA biosensor activity measured following expression of a panel of guanine nucleotide exchange factors (GEFs), n=3-11. Data represent mean ± SEM. Statistical significance was determined using an unpaired two-tailed Student’s t-test. *p<0.05, **p<0.01, ***p<0.001, ****p<0.0001. (G) RhoA biosensor GST-RBD-pulldown experiment. LinXe cells were transfected with the following: Lanes 1, untransfected control; 2, mCherry-RhoA F30L; 3, mCherry-RhoA T19N; 4, full length biosensor with F30L mutant RhoA; 5, full length biosensor with F30L mutant RhoA and PBD domain exchanged for the RBD domain; and 6, truncated biosensor with F30L mutant RhoA but without RBD.

**Supplemental Fig. S4 : Development and validation of a genetically encoded cyan-yellow Rap1B FRET biosensor**

Schematic representation of the Rap1B single-chain FRET biosensor. The biosensor consists of fluorescent proteins mCerulean3 and mVenus, flanked by C-terminally attached, full length Rap1B and the N-terminal Ras association domain (RBD) of RalGDS. (B) Representative emission spectra following donor excitation in cells expressing constitutively active (G12V) or inactive (S17N) Rap1B mutant biosensor. (C) Quantification of Rap1B biosensor activity measured in the biosensor constructs carrying mutations affecting Rap1B activation or the effector-binding interface, n=3-6. (D) Comparison of Rap1B biosensor activity in adherent and suspended cell conditions, n=5-6. (E) Rap1B biosensor activity measured following coexpression of Ras- and Rap-family guanine nucleotide exchange factors (GEFs), n=4-9. (F) Rap1B biosensor activity measured following coexpression of Rab-family GEFs, n=4-9. (G) Rap1B biosensor activity measured following coexpression of Rho-family GEFs, n=3-9. (H) Rap1B biosensor activity measured following coexpression of Rap GTPase-activating proteins (GAPs), n=4-9. Data represent mean ± SEM. Statistical significance was determined using an unpaired two-tailed Student’s t-test. *p<0.05, **p<0.01, ***p<0.001, ****p<0.0001. (I) Rap1B biosensor GST-RBD-pulldown experiment. LinXe cells were transfected with the following: Lanes 1, untransfected control; 2, mGreenLantern (mGL)-Rap1B Q63E; 3, mGL-Rap1B S17N; 4, full length biosensor with Q63E mutant Rap1B; and 5, full length biosensor with Q63E mutant Rap1B and PBD domain exchanged for the RBD domain.

**Supplemental Fig. S5: Signaling microdomains basic characterizations** (A) Average number of MDs / cell. (B) MD percent cell area occupancy. (C) Mean MD lifetimes. Data represent mean ± SEM. Statistical significance was determined using unpaired two-tailed Student’s t-tests. Pooled datasets from n=3 independent experiments.

**Supplemental Fig. S6: Uncropped Westernblots of Rap1A and RhoA biosensor stable cell line with or without induction**

**Supplementary Movie1: Representative Rap1A and RhoA biosensor activities imaged in a single living cell**

Representative time-series images of the Rap1A and RhoA biosensor activities for Control Rap1A (TOP) and RhoA (BOTTOM), shown along with DIC channel images. Images were taken at 10 s intervals. Frame rate 7 fps. White bar = 10 µm.

**Supplementary Movie2: Representative Rap1A and RhoA signaling MD analysis under control conditions**

Representative time-series images of the biosensor activities, signaling coordination strength, MDs, and biosensor activity-MD overlay are shown, for Control Rap1A (TOP) and RhoA (BOTTOM). Images were taken at 10 s intervals. Frame rate 7 fps. White bar = 10 µm.

**Supplementary Movie3: Representative Rap1A and RhoA signaling MD analysis under siARHGAP20 conditions**

Representative time-series images of the biosensor activities, signaling coordination strength, MDs, and biosensor activity-MD overlay are shown, for siARHGAP20 Rap1A (TOP) and RhoA (BOTTOM). Images were taken at 10 s intervals. Frame rate 7 fps. White bar = 10 µm.

**Supplementary Data1: Base-pair sequence of Rap1A biosensor**

**Supplementary Data2: Base-pair sequence of Rap1B biosensor**

**Supplementary Data3: Base-pair sequence of NIR RhoA biosensor**

## Author Contributions

C.S., S.P., N.C.C., R.B., R.R., D.S.V. and L.H. performed the experiments. R.R. wrote and optimized the computational pipelines. A.M. provided reagents, cDNA, and Rap1 biological guidance. G.D. supervised the computational analysis. J.A.A.-G. provided database for genetic screens and expression analysis guidance. C.S., S.P. and L.H. conceived the project. C.S., S.P., R.R. and L.H. reviewed the results and analysis. C.S., S.P. and L.H. wrote and revised the manuscript. All authors have read and agreed to the published version of the manuscript.

## Institutional Review Board Statement

Not applicable for studies not involving humans or animals. Document of Registration #250032 for Recombinant DNA, Albert Einstein College of Medicine, Department of Environmental Health and Safety.

## Informed Consent Statement

Not applicable for studies not involving humans.

## Data Availability Statement

The raw data supporting the conclusions of this article will be made available by the authors on request. The biosensor expression constructs are available from the authors on request. The image processing tools and computational algorithms are available from the authors on request.

## Acknowledgments

This work was supported by NIH grants R35GM136226 (L.H.), RM1GM145399 (G.D.), and the Chan Zuckerberg Initiative (L.H.). L.H. is an Irma T. Hirschl Career Scientist. J.A.A.-G. is supported by grants from The National Institute of Health (NIH) /National Cancer Institute (NCI) (CA109182, CA284085, CA301643, CA253977 and CCSG P30 CA013330-52), and The Gurwin Foundation and the Garfunkel, Spiegel and Spatz families for their generous support. J.A.A.-G. is also supported by a DOD CDMRP FY23 BCRP FL2 PSPS (BC230537-HT94252410078) and an Aging and Cancer grant from the Samuel Waxman Cancer Research Foundation and The Mark Foundation for Cancer Research. We thank members of the Segall and Cox laboratories at Albert Einstein College of Medicine for their helpful discussions.

