## Supplementary figures and images for "ARHGAP20 organizes spatial Rap1–RhoA signaling coordination controlling adhesion dynamics during migration"

### Supplementary Figure S1

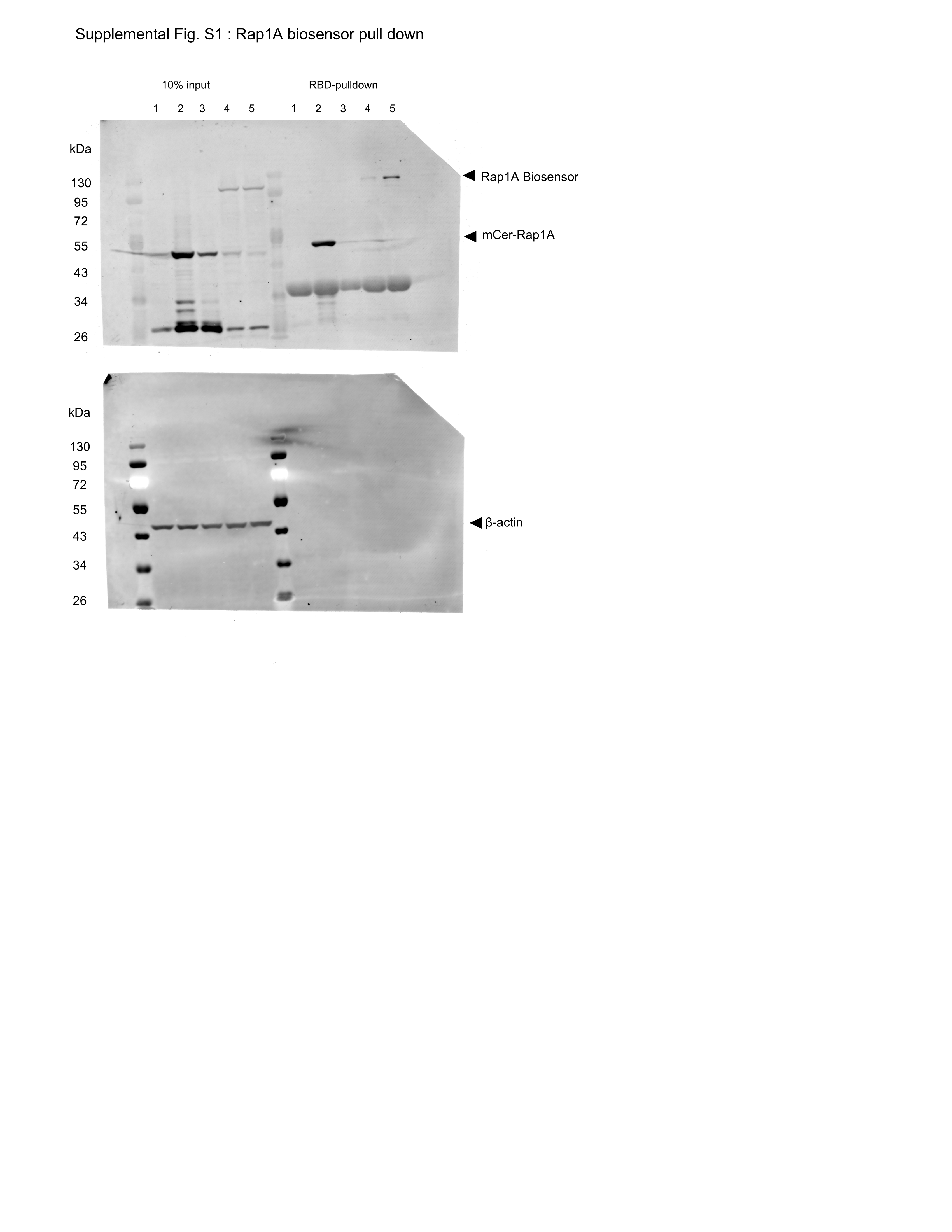

### Supplementary Figure S2

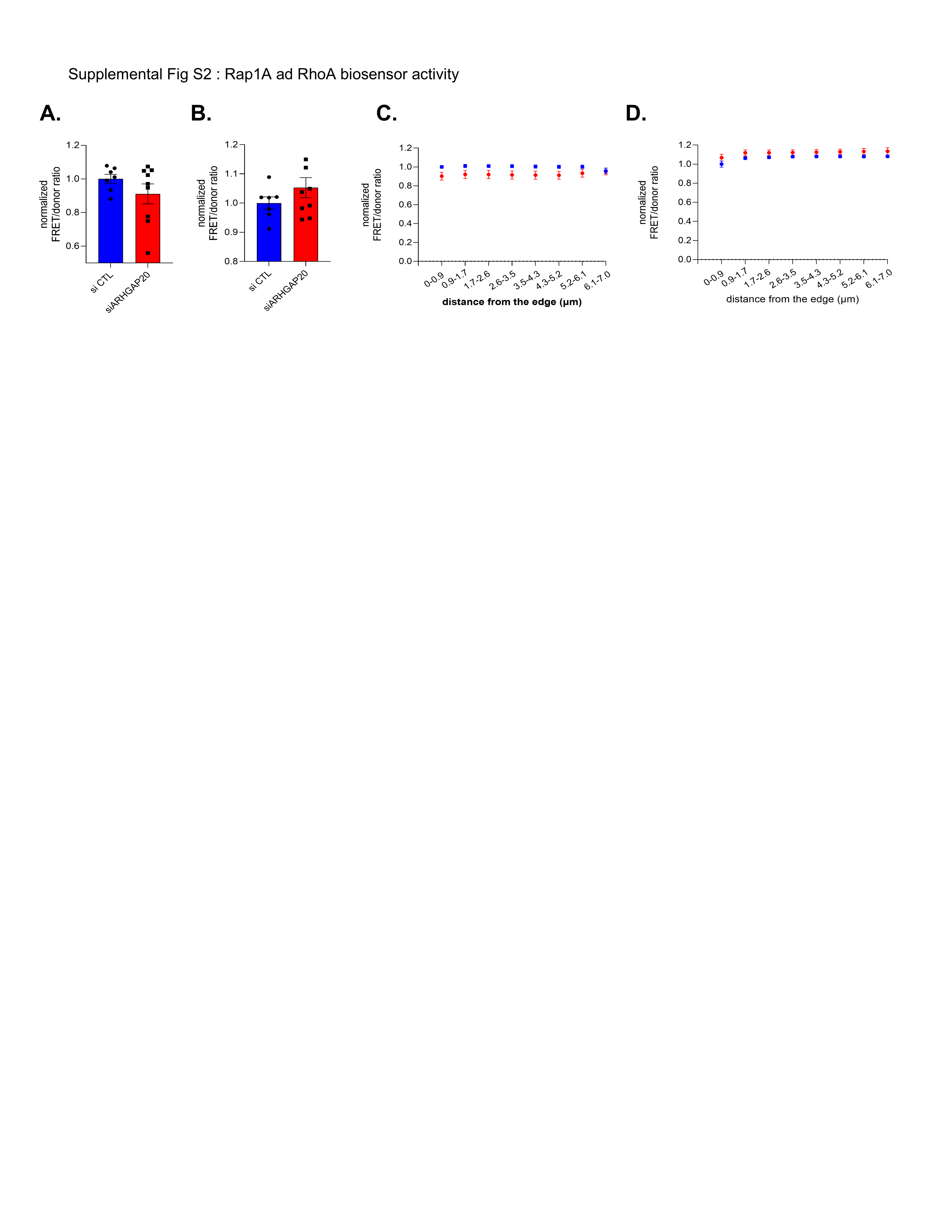

### Supplementary Figure S3

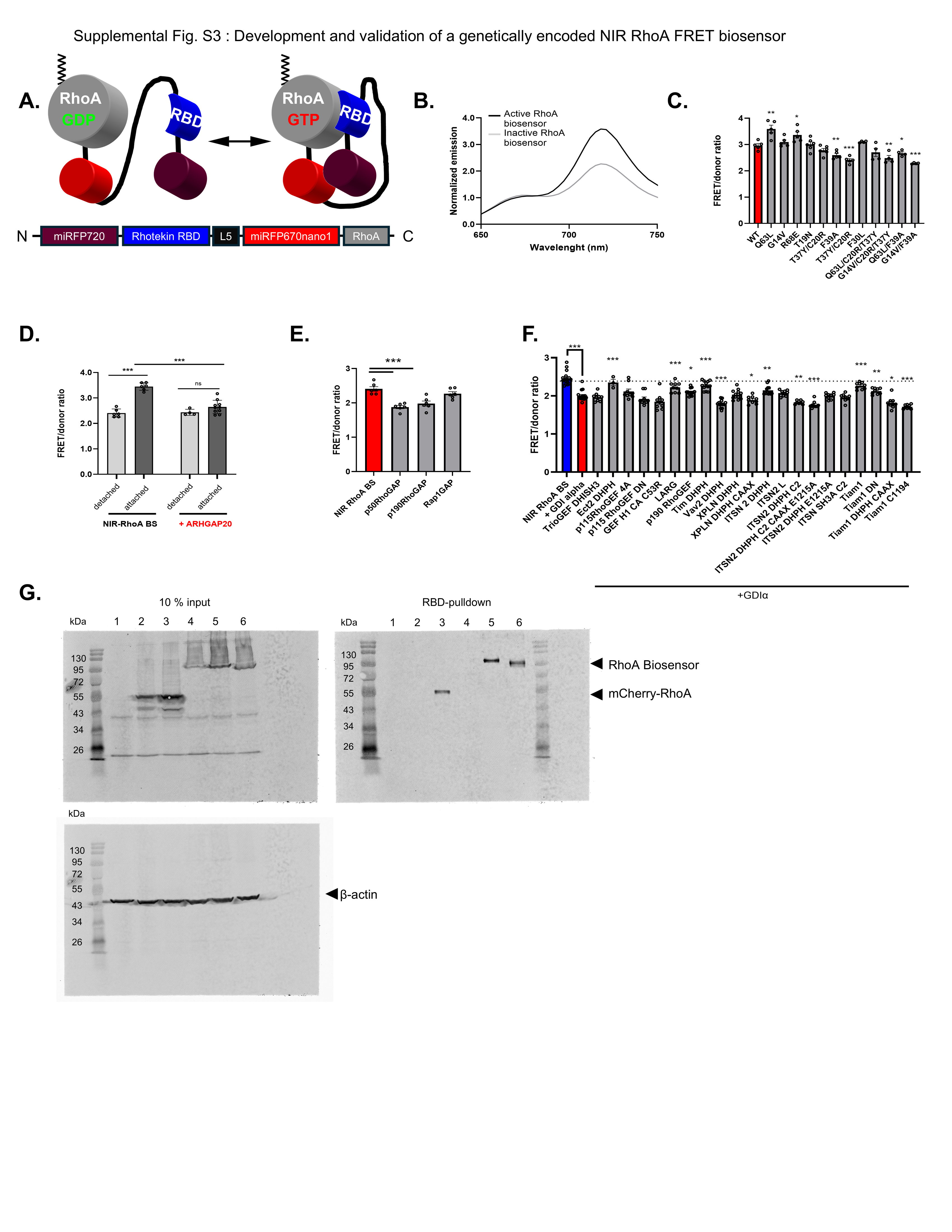

### Supplementary Figure S4

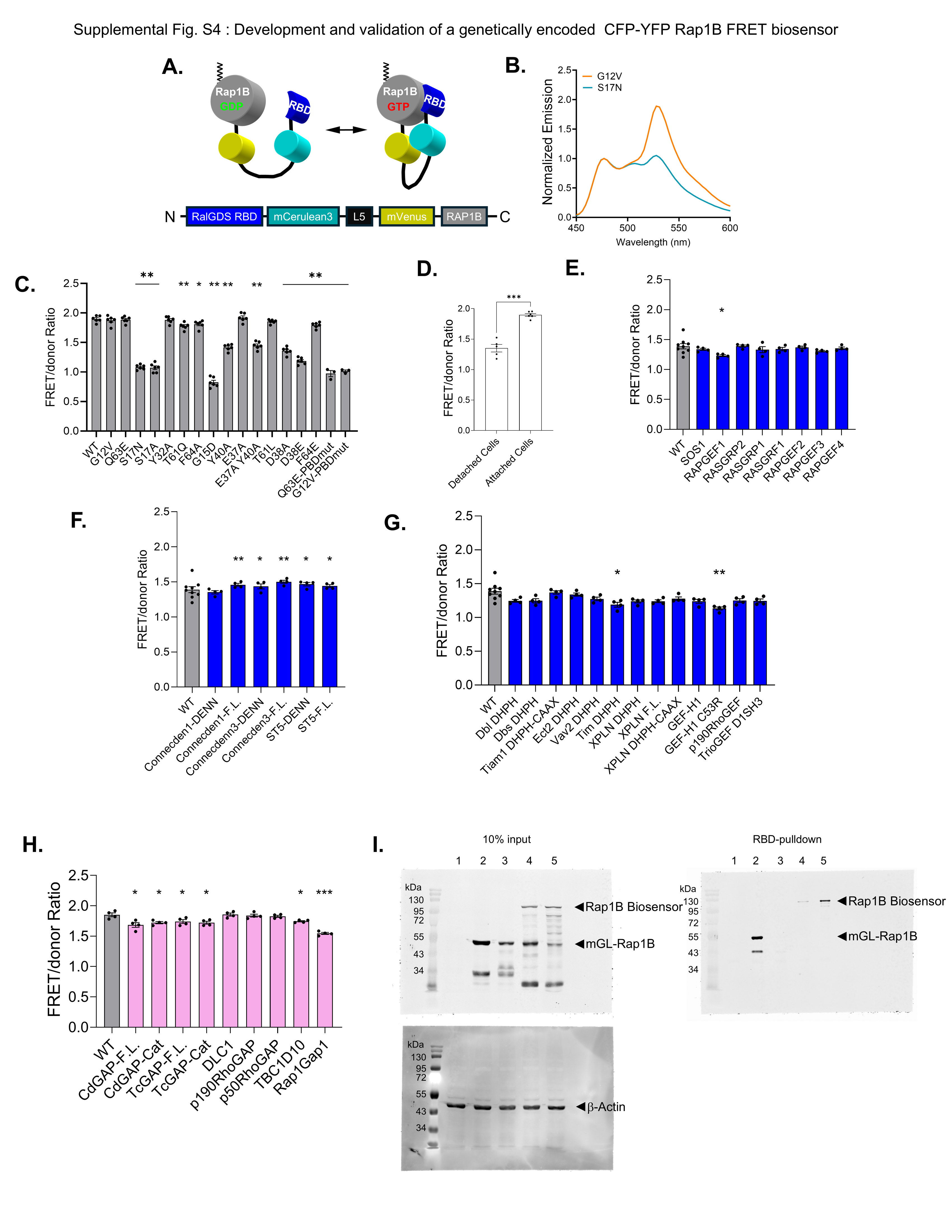

### Supplementary Figure S5

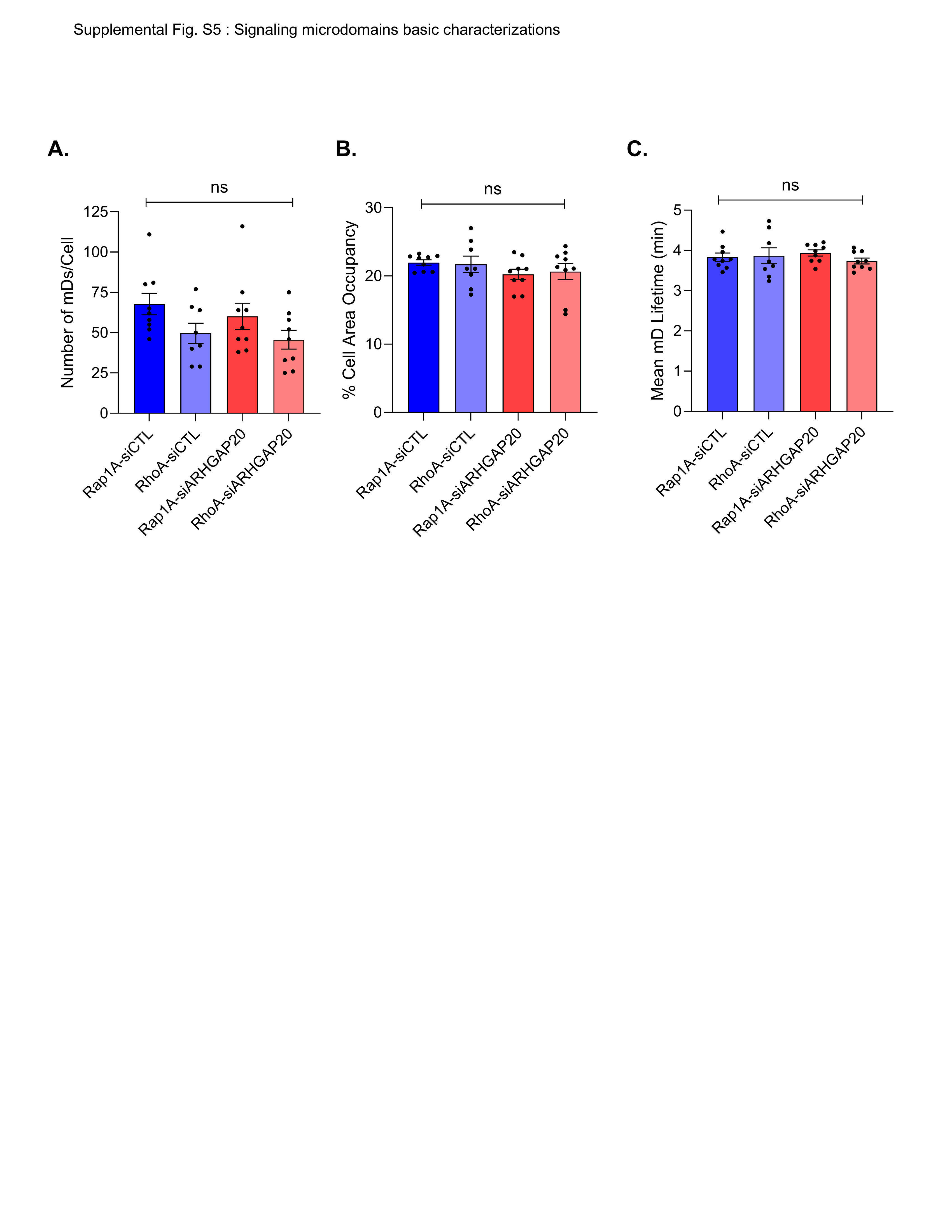

### Supplementary Figure S6

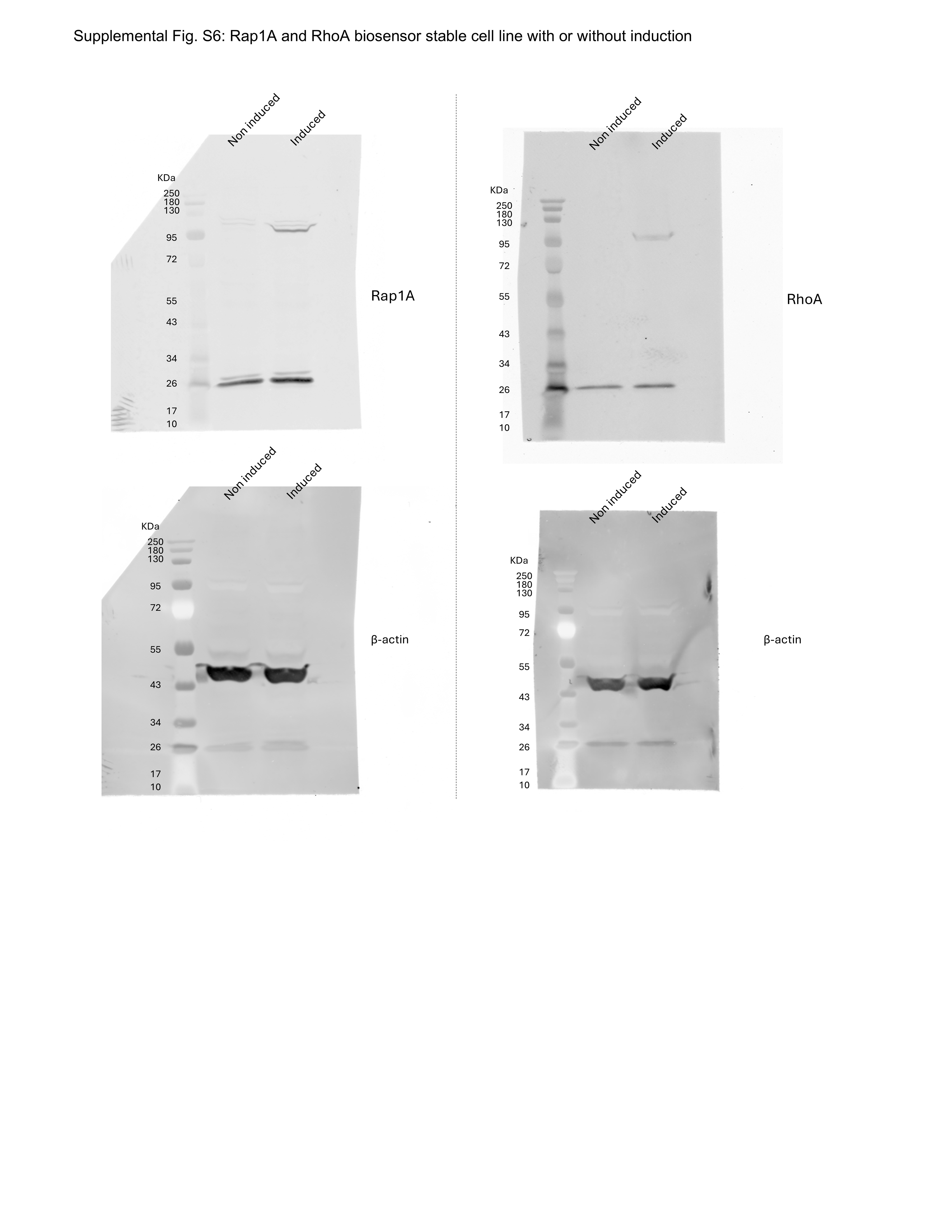
