## Supplementary Data1 for "ARHGAP20 organizes spatial Rap1–RhoA signaling coordination controlling adhesion dynamics during migration"

**Supplementary Data 1: Rap1A biosensor**

NcoI:

-CCatgG-CC

RalGDS-RBD:

tccgcgctgccgctctacaaccagcaggtgggcgactgctgtatcatccgcgtcagcctggacgtggacaatggcaacatgtacaagagcatcctggtgaccagccaagataaggctccggctgtaatccgcaaggcTatggacaaacacaacctggaggaggaggagccggaggactatgagctgctgcagattctctcagatgaccggaagctgaagatccctgaaaacgccaacgtcttctatgccatgaactctactgccaactatgactttgtcctgaagaagcggacc

BamHI:

-GGATCC-

mCerulean3-SM2:

ATGGTGTCCAAAGGAGAAGAACTGTTTACAGGAGTGGTCCCTATTCTGGTGGAACTGGATGGAGATGTGAATGGACATAAATTTTCCGTGAGCGGAGAAGGAGAAGGAGACGCTACATATGGAAAACTGACACTGAAGTTTATTTGTACAACAGGAAAACTGCCTGTGCCTTGGCCTACACTGGTGACCACACTGAGCTGGGGAGTCCAGTGTTTTGCTAGGTATCCTGATCATATGAAACAGCATGATTTCTTTAAAAGCGCTATGCCTGAGGGATATGTGCAGGAAAGGACAATTTTCTTTAAAGATGATGGAAATTATAAAACAAGGGCTGAAGTGAAATTTGAAGGAGATACACTGGTGAATAGGATTGAACTGAAAGGAATTGATTTTAAAGAAGATGGAAATATTCTGGGACATAAACTGGAATATAATGCTATTCACGGCAATGTGTACATTACAGCTGATAAACAGAAAAATGGAATTAAGGCTAATTTTGGCCTGAACTGCAATATTGAAGATGGAAGCGTGCAGCTGGCTGATCATTATCAGCAGAATACACCTATTGGAGATGGACCTGTGCTGCTGCCTGATAATCATTATCTGTCCACACAGAGCAAACTGTCCAAAGATCCTAATGAAAAAAGGGACCATATGGTGCTGCTGGAATTTGTGACAGCTGCCGGCATTACCCTGGGAATGGATGAACTGTATAA

HindIII:

-A-AGCTT-G

5-Linker SM2:

GCGGCCGGCACCAGCGGGTCTGGCAAAGGGTCTGGGGAGGGGAGCACCAAGGGAGGCAGCACGAGCGGCAGCGGGAAGGGAAGCGGCGAGGGATCTACGAAAGGGGGGAGCGGCACCTCCGGAAGCGGAAAGGGCAGCGGCGAGGGCAGCACGAAAGGTGGAAGCGGGACGTCCGGGTCCGGGAAAGGCTCCGGAGAGGGGTCCACAAAGGGCGGCTCCACTAGCGGCTCCGGCAAGGGGAGCGGGGAGGGCAGCACCAAGGGCGGCTCT

NotI:

-GCGGCCGC-A-

mVenus:

ATGGTGAGCAAGGGCGAGGAGCTGTTCACCGGGGTGGTGCCCATCCTGGTCGAGCTGGACGGCGACGTAAACGGCCACAAGTTCAGCGTGTCCGGCGAGGGCGAGGGCGATGCCACCTACGGCAAGCTGACCCTGAAGCTCATCTGCACCACCGGCAAGCTGCCCGTGCCCTGGCCCACCCTCGTGACCACCCTCGGCTACGGCCTGCAGTGCTTCGCCCGCTACCCCGACCACATGAAGCAGCACGACTTCTTCAAGTCCGCCATGCCCGAAGGCTACGTCCAGGAGCGCACCATCTTCTTCAAGGACGACGGCAACTACAAGACCCGCGCCGAGGTGAAGTTCGAGGGCGACACCCTGGTGAACCGCATCGAGCTGAAGGGCATCGACTTCAAGGAGGACGGCAACATCCTGGGGCACAAGCTGGAGTACAACTACAACAGCCACAACGTCTATATCACCGCCGACAAGCAGAAGAACGGCATCAAGGCCAACTTCAAGATCCGCCACAACATCGAGGACGGCGGCGTGCAGCTCGCCGACCACTACCAGCAGAACAC

CCCCATCGGCGACGGCCCCGTGCTGCTGCCCGACAACCACTACCTGAGCTACCAGTCCAAGCTGAGCAAAGACCCCAACGAGAAGCGCGATCACATGGTCCTGCTGGAGTTCGTGACCGCCGCCGGGATCACTCTCGGCATGGACGAGCTGTACAAG

EcoRI:

-GAATTC-

Rap1A (human)-WT:

ATGCGTGAGTACAAGCTAGTGGTCCTTGGTTCAGGAGGCGTTGGGAAGTCTGCTCTGACAGTTCAGTTTGTTCAGGGAATTTTTGTTGAAAAATATGACCCAACGATAGAAGATTCCTACAGAAAGCAAGTTGAAGTCGATTGCCAACAGTGTATGCTCGAAATCCTGGATACTGCAGGGACAGAGCAATTTACAGCAATGAGGGATTTGTATATGAAGAACGGCCAAGGTTTTGCACTAGTATATTCTATTACAGCTCAGTCCACGTTTAACGACTTACAGGACCTGAGGGAACAGATTTTACGGGTTAAGGACACGGAAGATGTTCCAATGATTTTGGTTGGCAATAAATGTGACCTGGAAGATGAGCGAGTAGTTGGCAAAGAGCAGGGCCAGAATTTAGCAAGACAGTGGTGTAACTGTGCCTTTTTAGAATCTTCTGCAAAGTCAAAGATCAATGTTAATGAGATATTTTATGACCTGGTCAGACAGATAAATAGGAAAACACCAGTGGAAAAGAAGAAGCCTAAAAAGAAATCATGTCTGCTGCTC-TAG

XhoI

-CTCGAG-
