## Supplementary Data3 for "ARHGAP20 organizes spatial Rap1–RhoA signaling coordination controlling adhesion dynamics during migration"

**Supplementary Data 3: NIR RhoA biosensor**

CC-

miRFP720-OPT:

atggcCgaGggCAGcgtGgccCGCcagccCgacctGCTGacctgcgacgaCgagCCCatccaCatccccggCgccatccaGccCcaCggCctgctgctGgccctGgccgccgacatgacCatcgtGgccggcagcgacaacctGcccgaGctGaccggCctggcCatcggcgccctgatcggccgcAGCgcCgccgaCgtGttcgacAGCgagacCcacaaccgCctgacCatcgccCTGgccgagcccggCgcCgccgtGggCgcCccCatcacCgtGggcttcacCatgcgCaaggacgcCggcttcatcggcAGCtggcaCcgccaCgaCcagctGatcttcctGgagctGgagccCccccagcgCgacgtGgccgagccCcaggcCttcttccgccgcaccaacagcgccatccgccgcctgcaggccgccgaGaccCTGgaGagcgcctgcgccgccgcCgcCcaGgaggtgcgCaagatCaccggcttcgaCcgCgtgatgatctaCcgcttcgccAGCgacttcagcggCAGCgtgatcgcCgaggaCcgCtgcgccgaggtGgagAGCaaGctGggcctgcactaCccCgccAGCttcatcccCgcCcaggcccgCcgCctGtaCaccatcaacccCgtGcgCatcatCcccgaCatcaaCtaCcgCccCgtgccCgtGaccccCgacctGaaCccCgtGaccggCcgCccCatCgaCctGagcttcgccatcctgcgcagcgtGAGCcccaaccaCctggagttcatgcgcaacatCggcatgcacggcacCatgAGCatcAGCatCCTGcgcggcgagcgCctgtggggCCTGatcgtGtgccaCcaccgCacCccCtactacgtGgaCctGgaCggccgccaGgcctgcaagCGCgtGgccgagCGCctggccacCcagatcggcgtgatggaGgag

BamHI:

-GGATCC-

RBD-SM1:

ATCCTGGAGGACCTgAAcATGCTgTACATCCGcCAGATGGCcCTgAGCCTGGAGGACACcGAGCTGCAGcgcAAgCTgGAcCAcGAGATCCGcATGcgcGAcGGcGCCTGCAAGCTGCTGGCcGCCTGCagcCAGCGcGAGCAGGCcCTGGAgGCCACCAAGAGCCTGCTGGTGTGCAACAGCCGcATcCTgAGCTACATGGGcGAGCTGCAGCGcCGcAAGGAGGCCCAGGTGCTGGAGAAGACc

Linker1-PP with HindIII:

GGCAGCGGCGGCAGCGGACCCCCCGGA-AGCTT-G

Linker2 with NotI:

ACTTCTGGTTCTGGTAAACCTGGTTCTGGTGAAGGTTCTACTAAAGGTGGATCT

ACaTCcGGaagcGGaAAACCaGGaTCtGGaGAgGGaagcACaAAAGGaGGgTCc

ACcagcGGcTCcGGcAAgCCcGGcagcGGcGAgGGcagcACcAAgGGcGGcagc

-GCGGCCGCA

miRFP670nano1:

atggcaaacctggacaagatgctgaataccacagtaacagaggtgcggcagttcctgcaggtggacagagtgtgcgtgttccagtttgaggaggattatagcggagtggtggtggtggaggccgtggacgataggtggatctccatcctgaagacccaggtgcgggatagatacttcatggagacaaggggcgaggagtattctcacggccgctaccaggccatcgccgacatctacaccgcaaacctgacagagtgctacagggatctgctgacacagtttcaggtgagagcaatcctggccgtgcccatcctgcagggcaagaagctgtggggcctgttggtggcacaccagctggcggcccctagacagtggcagacctgggagatcgactttctgaagcagcaggccgtggtggtgggcatcgccatccagcagagc

EcoRI:

-GAATTC-

RhoA-WT:

ATGGCTGCCATCCGGAAGAAACTGGTGATTGTTGGTGATGGAGCCTGTGGAAAGACATGCTTGCTCATAGTCTTCAGCAAGGACCAGTTCCCAGAGGTGTATGTGCCCACAGTGTTTGAGAACTATGTGGCAGATATCGAGGTGGATGGAAAGCAGGTAGAGTTGGCTTTGTGGGACACAGCTGGGCAGGAAGATTATGATCGCCTGAGGCCCCTCTCCTACCCAGATACCGATGTTATACTGATGTGTTTTTCCATCGACAGCCCTGATAGTTTAGAAAACATCCCAGAAAAGTGGACCCCAGAAGTCAAGCATTTCTGTCCCAACGTGCCCATCATCCTGGTTGGGAATAAGAAGGATCTTCGGAATGATGAGCACACAAGGCGGGAGCTAGCCAAGATGAAGCAGGAGCCGGTGAAACCTGAAGAAGGCAGAGATATGGCAAACAGGATTGGCGCTTTTGGGTACATGGAGTGTTCAGCAAAGACCAAAGATGGAGTGAGAGAGGTTTTTGAAATGGCTACGAGAGCTGCTCTGCAAGCTAGACGTGGGAAGAAAAA

ATCTGGTTGCCTTGTCTTGTGA

XhoI

-CTCGAG-
